# Beneficial Effects of Bempedoic Acid Treatment in Polycystic Kidney Disease Cells and Mice

**DOI:** 10.1101/2022.08.09.503392

**Authors:** Kenneth R. Hallows, Hui Li, Biagio Saitta, Saman Sepehr, Polly Huang, Jessica Pham, Jonathan Wang, Valeria Mancino, Eun Ji Chung, Stephen L. Pinkosky, Núria M. Pastor-Soler

## Abstract

ADPKD has few therapeutic options. Tolvaptan slows disease but has side effects limiting its tolerability. Bempedoic acid (BA), an ATP citrate-lyase (ACLY) inhibitor FDA-approved for hypercholesterolemia, catalyzes a key step in fatty acid/sterol synthesis important for cell proliferation. BA is activated by very long-chain acyl-CoA synthetase (FATP2) expressed primarily in kidney and liver. BA also activates AMPK. We hypothesized that BA could be a novel ADPKD therapy by inhibiting cyst growth, proliferation, injury, and metabolic dysregulation via ACLY inhibition and AMPK activation. *Pkd1*-null kidney cell lines derived from mouse proximal tubule (PT) and collecting duct (IMCD) were grown in 2D or 3D Matrigel cultures and treated ± BA, ± SB-204990 (another ACLY inhibitor) or with *Acly* shRNA before cyst analysis, immunoblotting or mitochondrial assays using MitoSox and MitoTracker staining. *Pkd1*^*fl/fl*^*;Pax8-rtTA;Tet-O-Cre* C57BL/6J mice were induced with doxycycline injection on postnatal days 10 and 11 (P10-P11) and then treated ± BA (30 mg/kg/d) ± tolvaptan (30-100 mg/kg/d) by gavage from P12-21. Disease severity was determined by % total-kidney-weight-to-bodyweight (%TKW/BW) and BUN levels at euthanasia (P22). Kidney and liver homogenates were immunoblotted for expression of key biomarkers. ACLY expression and activity were upregulated in *Pkd1*-null PT and IMCD-derived cells vs. controls. Relative to controls, both BA and SB-204990 inhibited cystic growth in *Pkd1*-null kidney cells, as did *Acly* knockdown. BA inhibited mitochondrial superoxide production and promoted mitochondrial elongation, suggesting improved mitochondrial function. In ADPKD mice, BA reduced %TKW/BW and BUN to a similar extent as tolvaptan vs. untreated controls. Addition of BA to tolvaptan caused a further reduction in %TKW/BW and BUN vs. tolvaptan alone. BA generally reduced ACLY and stimulated AMPK activity in kidneys and livers vs. controls. BA also inhibited mTOR and ERK signaling and reduced kidney injury markers. In liver, BA treatment, both alone and together with tolvaptan, increased mitochondrial biogenesis while inhibiting apoptosis. We conclude that BA and ACLY inhibition inhibited cyst growth in vitro, and BA decreased ADPKD severity in vivo. Combining BA with tolvaptan further improved various ADPKD disease parameters. Repurposing BA may be a promising new ADPKD therapy, having beneficial effects alone and along with tolvaptan.

## INTRODUCTION

Autosomal dominant polycystic kidney disease (ADPKD), the most common genetic cause of end-stage kidney disease (ESKD), affects every ethnicity with a prevalence of ∼1:500-1000 and ∼600,000 patients in the U.S. alone (Gabow, 1993, Chebib and Torres, 2016). Patients with ADPKD present with enlarging cystic lesions in the kidney and often the liver as well, leading to a progressive decline in kidney function that is associated with ESKD in half of ADPKD patients by age 50-60 (Chebib and Torres, 2016). Most ADPKD patients have loss-of-function mutations in the multifunctional proteins polycystin-1 or -2 (PC1 and PC2, encoded by the genes *PKD1* and *PKD2*) (Chebib and Torres, 2016). ADPKD therapeutic options to specifically address the decline in glomerular filtration rate (GFR) are very limited. The only current FDA-approved drug for ADPKD is tolvaptan, a vasopressin 2 receptor (V2R) antagonist. This drug slows disease progression in patients at risk for rapid progression towards end-stage kidney disease (ESKD) (Torres et al., 2012). However, tolvaptan has the dose-dependent side effect of polyuria and a risk of hepatotoxicity that requires monthly monitoring of liver function tests (Blair, 2019, Woodhead et al., 2017). Thus, there is a clear need for additional ADPKD therapies targeting different cellular pathways dysregulated in ADPKD that could potentially be used alone or in combination with tolvaptan.

There is growing recognition that ADPKD cyst-forming PC1-deficient cells have major metabolic derangements that likely contribute to kidney tubular epithelial cyst formation and expansion. Specifically, compared to control kidney tubular epithelial cells, ADPKD cells display increased aerobic glycolysis (the Warburg effect), impaired fatty acid oxidation, increased cellular proliferation, and reduced AMP-activated protein kinase (AMPK) activity (Rowe et al., 2013, Menezes et al., 2016). Earlier, we helped pioneer the use of the AMPK activator metformin to inhibit ADPKD kidney cyst growth in mouse models of *Pkd1* knockout (Takiar et al., 2011).

AMPK is a ubiquitous metabolic sensor that regulates many cellular processes (Hallows, 2005, Steinberg and Kemp, 2009, Steinberg and Carling, 2019). The role of AMPK in the protection of kidney function has been studied in many models of acute and chronic kidney disease (Rajani et al., 2017). Of note, kidney AMPK activity is generally decreased in both humans and mice with chronic kidney disease (CKD) (Dugan et al., 2013, Li et al., 2015). The renoprotective role of AMPK in CKD is thought to occur through activation and induction of several effector pathways including autophagy, fatty acid oxidation, antioxidant pathways (Decleves et al., 2011) and via inhibition of the inflammatory cascade (Peairs et al., 2009). In response to metabolic and other cellular stresses, AMPK activation helps maintain cellular energy balance by restoring ATP levels through regulation of metabolic enzymes, promoting cellular energy efficiency, and inhibiting pro-growth anabolic pathways.

Our group recently demonstrated that the AMPK activator metformin ameliorates relevant disease parameters in a hypomorphic PKD mouse model that closely mimics human ADPKD (Pastor-Soler et al., 2022). We have also been involved in the TAME-PKD study where metformin was found to be safe and tolerable in ADPKD patients (Perrone et al., 2021). Of note, metformin doses that inhibit cyst growth in pre-clinical ADPKD models may not be as tolerable or clearly efficacious in ADPKD patients (Brosnahan et al., 2022, Perrone et al., 2021). Moreover, along with the various beneficial effects of metformin, including inhibition of cyst fluid secretion, cell proliferation and cAMP production (Takiar et al., 2011, Miller et al., 2013), metformin inhibits Complex I of the mitochondrial respiratory chain (Owen et al., 2000), which may hamper the promotion of defective mitochondrial oxidative metabolism in ADPKD. Thus, novel drugs targeting complementary pathways in ADPKD that could potentially synergize with tolvaptan or metformin may afford lower effective drug dosing and have better efficacy against the disease when used in combination in patients.

Here we explored targeting and inhibiting the enzyme ATP-citrate lyase (ACLY) to determine its effects on relevant disease parameters in vitro and in a conditional *Pkd1* knockout mouse model of ADPKD. ACLY is a key metabolic enzyme that promotes lipid and cholesterol biosynthesis by generating acetyl-CoA from cytosolic citrate and has been recently identified as a new therapeutic target for lowering cholesterol in patients with atherosclerotic cardiovascular disease (Huynh, 2019). In addition, ACLY inhibition has emerged as a new therapeutic strategy for cancer, where ACLY inhibition blocks lipid synthesis and cellular proliferation (Zhao et al., 2016). As with tumor growth, cystic growth in ADPKD relies on such mechanisms that support enhanced cellular proliferation.

Importantly, ACLY has also been reported to bind to and inhibit the AMPK-β _1_ subunit (Lee et al., 2015), suggesting a mutual antagonism between ACLY and AMPK.

Bempedoic acid (BA; also known as ETC-1002) inhibits ACLY and is approved by the FDA as an adjunct to diet and maximally tolerated statin therapy for the treatment of adults with atherosclerotic cardiovascular disease who require additional lowering of LDL-C. BA also activates AMPK in mice (Pinkosky et al., 2016). BA exists as a pro-drug that gets converted to its active form by an enzyme (Very long-chain acyl-CoA synthetase; ACSVL1 or FATP2) whose tissue expression is primarily limited to kidney and liver (Pinkosky et al., 2016), the two principal organs affected in ADPKD. We thus hypothesized that BA treatment could be beneficial in ADPKD by inhibiting kidney cyst growth, inflammation, injury, and metabolic dysregulation via simultaneous ACLY inhibition and AMPK activation with limited off-target effects. The rationale for the use of BA in ADPKD involves correcting the dysregulated metabolism and excessive cell proliferation in ADPKD and is summarized schematically in **Figure 1**. Herein, we tested the potential beneficial effects of BA alone and in combination with tolvaptan on 3D cyst growth and mitochondrial function in *Pkd1*-null kidney cells in vitro, and on key parameters of disease severity in kidneys and liver in conditional *Pkd1* knockout mice.

**Figure 1.**
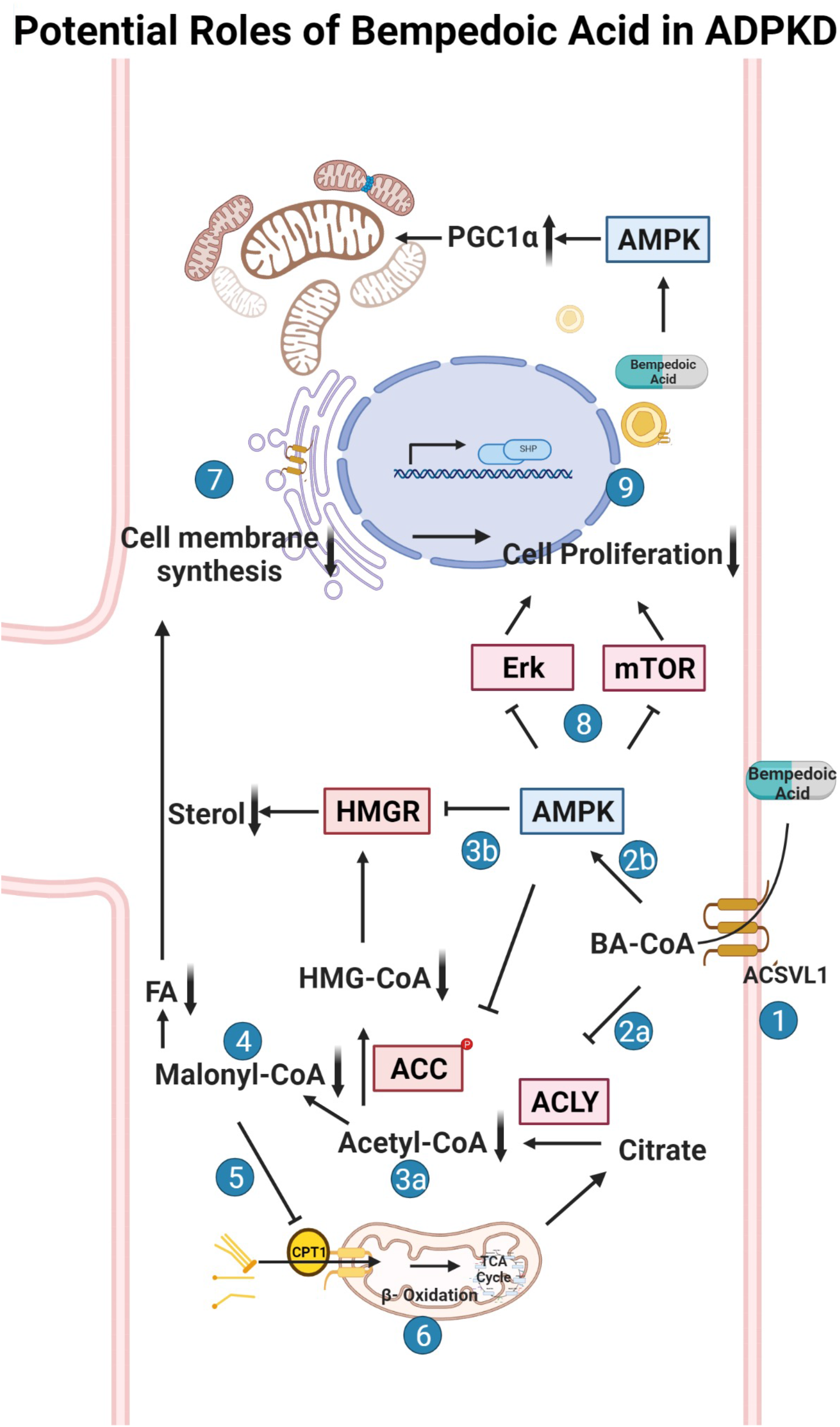
Potential roles of bempedoic acid in ADPKD. Schematic flow diagram of the effects of bempedoic acid (BA; a.k.a. ETC-1002) on various metabolic pathways and cellular proliferation via inhibition of ATP-citrate lyase (ACLY) and activation of AMP-activated protein kinase (AMPK). 1. The pro-drug BA gets converted to its active form (BA-CoA) via Very Long-Chain Acyl-CoA Synthetase (ACSVL1; a.k.a. FATP2), whose long isoform is only substantially expressed in liver and kidney tissues (Pinkosky et al., 2016, Steinberg et al., 1999). BA-CoA simultaneously inhibits ACLY (2a) and activates AMPK (2b) in cells. Inhibition of ACLY, which converts citrate to acetyl-CoA and oxaloacetate, results in decreased cytosolic acetyl-CoA production (3a). Activated AMPK inhibits HMG-CoA reductase (HMGR) and acetyl-CoA carboxylase (ACC) (3b), which along with decreased acetyl-CoA production, inhibits the formation of malonyl-CoA (4) and HMG-CoA and thus inhibits both sterol and fatty acid (FA) synthesis. Decreased levels of malonyl-CoA, which is an inhibitor of carnitine palmitoyltransferase-1 (CPT-1), promotes FA uptake into mitochondria via CPT-1 (5) and thus promotes FA beta oxidation in mitochondria via the tricarboxylic acid (TCA) cycle (6). Decreased sterol and FA synthesis inhibits the synthesis of cellular membranes (7), which along with AMPK-dependent inhibition of various cellular pathways (incl. the mammalian target of rapamycin (mTOR) pathway) (8), causes inhibition of cellular proliferation and thus inhibition of cyst growth and expansion in ADPKD (9).

## MATERIALS AND METHODS

### Reagents and chemicals

All reagents and chemicals used were purchased from Sigma (St. Louis, MO, USA) or Thermo Fisher (Pittsburgh, PA, USA) unless otherwise stated. Pharmaceutical grade bempedoic acid (BA) and tolvaptan were obtained from Esperion Therapeutics, Inc. (Ann Arbor, MI, USA) and Otsuka Pharmaceuticals (Japan), respectively. Please see **Table 1** for detailed information on the antibodies and conditions that were used for immunoblotting.

**Table 1.** Antibodies Used for Western Blot.

| Antigen | Manufacturer | Cat. # | Host | Dilution | Incubation |  |
| --- | --- | --- | --- | --- | --- | --- |
|  |  |  |  |  | Time | Temperature |
| p-ACC(Ser79) | Cell Signaling | 3661 | Rabbit polyclonal | 1:1,000 | O/N | 4°C |
| p-ACLY(Ser455) | Cell Signaling | 4331 | Rabbit polyclonal | 1:1,000 | O/N | 4°C |
| t-ACLY | Cell Signaling | 4332 | Rabbit polyclonal | 1:1,000 | O/N | 4°C |
| p-AMPKα(Thr172) | Cell Signaling | 2531 | Rabbit polyclonal | 1:1,000 | O/N | 4°C |
| p-ERK(Thr202/Tyr204) | Cell Signaling | 9101 | Rabbit polyclonal | 1:1,000 | O/N | 4°C |
| Cleaved Cas3(Asp175) | Cell Signaling | 9661 | Rabbit polyclonal | 1:1,000 | O/N | 4°C |
| PGC-1α | Santa Cruz | SC-517380 | Mouse monoclonal | 1:500 | O/N | 4°C |
| pP70S6K(Thr389) | Santa Cruz | SC-11759R | Rabbit polyclonal | 1:500 | O/N | 4°C |
| NGAL | Abcam | ab63929 | Mouse polyclonal | 1:1,000 | O/N | 4°C |
| KIM-1 | R&D systems | AF1817 | Goat polyclonal | 1:800 | O/N | 4°C |
| FATP2 | Proteintech | 14048-1-AP | Rabbit polyclonal | 1:1000 | O/N | 4°C |
| Revert 700 total protein stain kit | LI-COR | 926-11010 |  |  | 5 min | RT |

O/N – Overnight; p – Phosphorylated or phosphor-; RT – Room Temperature; T – Total.

### Cell culture models

*Pkd1*-Null (PN24) and *Pkd1*-Het (PH2) cells were a kind gift of Dr. Stefan Somlo and were originally derived by microscopically dissecting and then dissociating proximal tubules (PTs) from *Pkd1*^flox/-^:TSLargeT (ImmortoMouse) mice. Parental *Pkd1*^flox/-^ cells from a clone showing epithelial properties were then transiently transfected with a plasmid encoding Cre recombinase and cloned again by limiting dilution, giving rise to daughter cells that either expressed Cre and therefore had undergone transformation to *Pkd1*^*-/-*^ or that had not expressed Cre and retained the parental *Pkd1*^flox/-^ genotype, as described previously (Joly et al., 2006). In this manuscript, we also refer to the PH2 cells as *Pkd1*^*+/-*^ cells for simplicity. These cells were cultured in medium composed of equal volumes of DMEM and Ham’s F-12 plus 7.5 nM sodium selenate, 5 μg/ml transferrin, 2 mM glutamine, 2 nM triiodothyronine, 5 μg/ml insulin, and 2% (vol/vol) FBS. Cells were maintained under permissive conditions (33°C with 10 U/ml γ-interferon) in a humidified 5% CO_2_-95% air incubator with medium changes every other day and passaged at least twice weekly. To induce differentiation, cells were kept under non-permissive conditions (37°C without γ-interferon) for 5-7 days prior to use in experiments. Mouse inner medullary collecting duct (IMCD) cell lines were also generously provided by Dr. Stefan Somlo. Wild-type (Wt)-IMCD3 (Cas9) is a control cell line for IMCD3-CRISPR-knockout cells (ID1-3E5 *Pkd1*^-/-^) (Decuypere et al., 2021). Both cell lines were cultured in regular DMEM/F12 medium containing 5% FBS in a 37°C humidified 5% CO_2_-95% air incubator with medium changes every other day and passaged approximately twice weekly.

### Generation of ACLY knockdown cell lines

A non-silencing lentiviral shRNA (pGIPZ) was used as a control and three different ACLY shRNA lentiviral constructs (pGIPZ) were obtained from Horizon Discovery (Waterbeach, Cambridge, UK). Recombinant lentiviral particles were produced by transient transfection of 293T cells according to the manufacturer’s protocol. The PT-derived *Pkd1*^*-/-*^ (PN24) and *Pkd1*^+/-^ (PH2) cells were infected with the cell culture supernatant containing lentiviral particles for 48 h. These cells were then selected in puromycin to generate stable cell lines with non-silencing and ACLY-specific shRNA. Cell lines were validated for diminished ACLY expression by Western blot analysis.

### 3D culture

Matrigel™ from BD Biosciences (BD #356234) was thawed overnight at 4°C prior to adding 50 μl to each well of an 8-well glass chamber slide (Lab-Tek #155409) and spreading evenly in the well using a P-200 tip. The slides were then placed in a cell culture incubator to allow the Matrigel™ to solidify for at least 15 minutes. During this time, PH2 *Pkd1*^*+/-*^ cells or PN24 or ID1-3E5 *Pkd1*^-/-^ cells were trypsinized and mixed into a stock of culture medium containing 2% Matrigel™ (assay medium) at a density of 6,000 cells per 400 μl of this medium. 400 μl of this mixture was then plated on top of the solidified Matrigel™ in each well of the chamber slide on day 0. Cells were then allowed to grow in a 5% CO_2_ humidified incubator at 37°C and fed with assay medium every other day for 12 days. Starting on day 1 after seeding, 3D cell cultures were treated with 10 µM forskolin plus 100 µM isobutylmethylxanthine (IBMX), in the absence or presence of 100 µM BA or 30 µM SB-204990 for the last 3 days of culture before imaging and cyst size analysis. DMSO was the vehicle control. For the studies shown in Fig. 5, PH2 or PN24 cells stably transduced to express either non-silencing or *Acly* shRNA constructs were used. For analysis, cysts (fluid-filled cell structures) were thresholded from background, and the cross-sectional area of each cyst grown in 3D Matrigel culture was calculated using the ImageJ Analyze Particles plug-in software (NIH).

### Mitochondrial morphology and superoxide quantification

To analyze mitochondrial morphology and superoxide production as indirect measures of mitochondrial health, PT- and IMCD-derived *Pkd1*^-/-^ cell lines were treated with vehicle or BA (100 µM) and then stained with MitoTracker™ Green FM (M7514), MitoTracker™ Deep Red FM (M22426) or MitoSOX™ Red mitochondrial superoxide indicator (M36008), along with the nuclear stain Hoechst 33342 (H3570; ThermoFisher Scientific Inc. Pittsburgh, PA, USA). Briefly, PN24 or ID1-3E5 cells were seeded onto 4-well chamber slides in the above-described media prewarmed to 37°C. Staining and washes were performed 2 days after plating and 24 h after the indicated treatments according to the manufacturer’s protocols. Cells were imaged using a Leica DMi8 live cell imaging fluorescence microscope using a 40X oil objective at zoom 1.6. Automated quantification of mitochondrial morphology was done on MitoTracker-stained cells using ImageJ software. The aspect ratio (length/width) was used as a measurement of mitochondrial elongation. We compiled the mean aspect ratio of >1,000 mitochondria in each of the selected cells and then compared the mean values (± SEM) from each of the cells analyzed across the two conditions (± BA treatment). MitoSOX Red fluorescence was used to assess mitochondrial superoxide production. The fluorescence intensity of each randomly selected cell was quantified using ImageJ software. Results are reported as compiled mean cellular values ± SEM from four independent biological replicate experiments for PN24 cells (with n = 159-188 cells analyzed) in **Figure 6** and one experiment for ID1-3E5 cells (n = 26-31 cells analyzed) in **Supplementary Fig. 2**.

### ADPKD mouse models

All animal procedures followed NIH guidelines for the care and use of laboratory animals and were approved by the University of Southern California’s Institutional Animal Care and Use Committee. Male and female *Pkd1*^*fl/fl*^*;Pax8-rtTA;Tet-O-Cre* transgenic mice in the C57BL/6J background were obtained as a generous gift from the Baltimore PKD Core Center and were used as an ADPKD model for in vivo studies, as described previously (Ma et al., 2013). Genotyping was confirmed between postnatal days 5 and 7 (P5-P7) and *Pkd1* inactivation was induced with IP doxycycline injection (50 mg/kg) on P10 and P11 to induce rapidly progressive cyst development. Mice were then treated in the absence or presence of BA (30 mg/kg/d) with or without cotreatment with tolvaptan (30-100 mg/kg/d) by daily oral gavage from P12-21. These drugs (or vehicle) were formulated into micelle suspensions comprised of 1,2-distearoyl-sn-glycero-3-phosphoethanolamine-N-methoxy-poly(ethylene glycol 2000) (DSPE-PEG(2000)-maleimide, Avanti Polar Lipids, Alabaster, AL, USA) to improve solubility prior to daily gavage injection by syringe into the mouse pups. Drug-loaded micelles were self-assembled via thin film evaporation using previously described methods (Huang et al., 2020). Briefly, 18 mg of BA (and/or up to 66 mg of tolvaptan) and 100 µM of DSPE-PEG(2000)-methoxy were dissolved in methanol and evaporated with nitrogen gas to form thin films. The resulting thin films were dried overnight under vacuum and hydrated in PBS at 80 °C for 30 min, and drug-loaded micelles were stored at 4°C and used within 3 days of formulation. Mice were euthanized at the end of the experiment at P22 to harvest kidneys and livers and obtain blood to assess various measures of disease severity.

### Whole blood chemistry measurements

We used the Abbott i-STAT handheld blood analyzer equipped with Chem 8+ cartridges for measurements blood urea nitrogen (BUN) from mixed venous blood at the time of euthanasia at P22 (Tinkey et al., 2006). Briefly, mice were anesthetized with isoflurane, and approximately 100 µl of blood was obtained from the submandibular venous plexus using a single-use lancet (Golde et al., 2005). The blood was collected in a heparinized vial and quickly added to the i-STAT cartridge, and results were obtained within two minutes.

### Kidney weight measurements, tissue preparation and microscopy

After clamping the renal pedicle at P22 the left kidney was quickly removed, rapidly weighed, sectioned coronally in two parts and placed in microcentrifuge tubes and frozen in liquid nitrogen. The right kidney was also removed after clamping of the renal pedicle, quickly weighed, and rapidly cut coronally. One half of this right kidney was fixed in 4% paraformaldehyde (Electron Microscopy Sciences, Hatfield, PA) in PBS, and the other half was placed in RNAlater (Qiagen, Waltham, MA) for future preparation of cDNA. A section of liver was also obtained for each mouse and placed immediately in liquid nitrogen.

After overnight fixation at 4°C, the kidney tissues were washed in PBS, quenched in NH_4_Cl and further washed in PBS. The samples were then placed in 10% neutral buffered formalin (VWR, Radnor, PA, USA) at 4°C for 16-18 hours. The fixed kidney samples were then dehydrated in serial alcohols, then cleared with xylene, embedded in paraffin, and cut into 4-μm sections on a rotary microtome at the Keck School of Medicine of USC Norris Pathology Core. The fixed tissues were stained with hematoxylin and eosin (H&E) for histological evaluation. We then obtained images using an Olympus IX73 inverted microscope using a Plan Achromat 2X objective for a total magnification of 20X (numerical aperture of 0.06 and working distance of 5.8mm).

### Electrophoresis and immunoblotting analysis

Kidney and liver lysates were prepared from frozen tissues, homogenized, centrifuged, proteins quantitated, and samples electrophoresed and transferred to nitrocellulose membranes, as described previously (Pastor-Soler et al., 2022). The membrane was first stained with Revert 700 total protein stain solution (LI-COR) and then washed prior to imaging at 700 nm for total protein quantitation per lane, as per the manufacturer’s recommendations. After destaining, the membrane was then blocked and probed with primary and secondary antibodies of interest. Quantification of immunoblot images was performed by densitometry and analyzed using Image Studio Lite Ver 5.2 software (LI-COR, USA). Please refer to Table 1 for information on antibodies and blotting conditions.

### Statistics

Statistical analysis was performed using GraphPad Prism (GraphPad, La Jolla, CA, USA) to obtain the mean values and SEM for each treatment group. In most experiments, significance was determined using two-tailed, unpaired Student’s t-tests assuming unequal variances for the groups, or one-way ANOVA with post-hoc Tukey corrections for multiple comparisons. Individual data points are shown in the bar graphs of figures, along with the mean (± S.E.) for each treatment condition. *P* values <0.05 were considered significant.

## RESULTS

As outlined above, the purpose of this initial study was to test as a proof of concept the potential role of the ACLY inhibitor and AMPK activator BA as a novel therapeutic for ADPKD using in vitro and in vivo ADPKD model systems. A particular feature of BA that may lend itself specifically to ADPKD therapy is its expected activity limitation primarily to kidney and liver, the two organs predominantly affected in the disease, where the BA pro-drug can get converted to its active metabolite via local expression of very long-chain acyl-CoA synthetase (FATP2 or ACSVL1; Cf. **Figure 1**, step 1), as described below.

### Very long-chain acyl-CoA synthetase is detected in mouse kidney, liver, and in key model cell lines for the study of ADPKD

To determine whether our kidney cell lines and conditional *Pkd1* KO mouse model would be useful tools to evaluate the potential beneficial effects of BA in reducing cyst size or number in ADPKD, we first evaluated FATP2 protein expression in cells and tissues by immunoblot. Specifically, two bands were detected using an anti-FATP2 primary antibody that detects both a shorter FATP2 splice variant (FATP2b) and the long form of FATP2a at ∼70kDa (Melton et al., 2011). The FATP2b variant lacks the protein domain required for the conversion of BA to its active metabolite (BA-CoA) and was expressed in all mouse tissues tested (**Figure 2A**, *lower band*). However, the FATP2a form was previously reported to have significant expression only in liver and kidney tissue, thus providing specificity of BA pro-drug conversion to its active form only in these organs (Pinkosky et al., 2016, Steinberg et al., 1999). Consistent with these earlier studies, we detected the full-length FATP2a primarily in liver and kidney tissue homogenates, with minimal expression in other tissues (**Figure 2A**, *upper panel, upper band*). Importantly, FATP2 immunoblotting of representative *Pkd1*^*fl/fl*^; *Pax8-rtTA; Tet-O-Cre* mouse kidney homogenates treated with or without doxycycline to induce tubule-specific *Pkd1* gene inactivation revealed expression of both FATP2 isoforms in both induced and uninduced mouse kidneys (**Figure 2B**). Quantification of the immunoblot showed that the kidneys of induced mice, the mice that developed ADPKD upon doxycycline injection, expressed less FATP2a than the uninduced mice. We also detected both FATP2a and FTAP2b isoforms in immortalized mouse kidney epithelial cells derived from both PT (**Figure 2C**) and IMCD (**Supplementary Figure 1**). Of note, densitometric quantification of the immunoblots revealed significant decreases in FATP2a expression in both *Pkd1-null* kidney tissue and cells compared to controls (**Figure 2** and **Supplementary Figure 1**). Altogether, these results confirm that the ADPKD cell lines and ADPKD mouse model used in our studies express the enzyme required to convert the BA pro-drug to its active compound and help support a rationale to evaluate the effects of BA on cyst growth in cell culture and in vivo.

**Figure 2.**
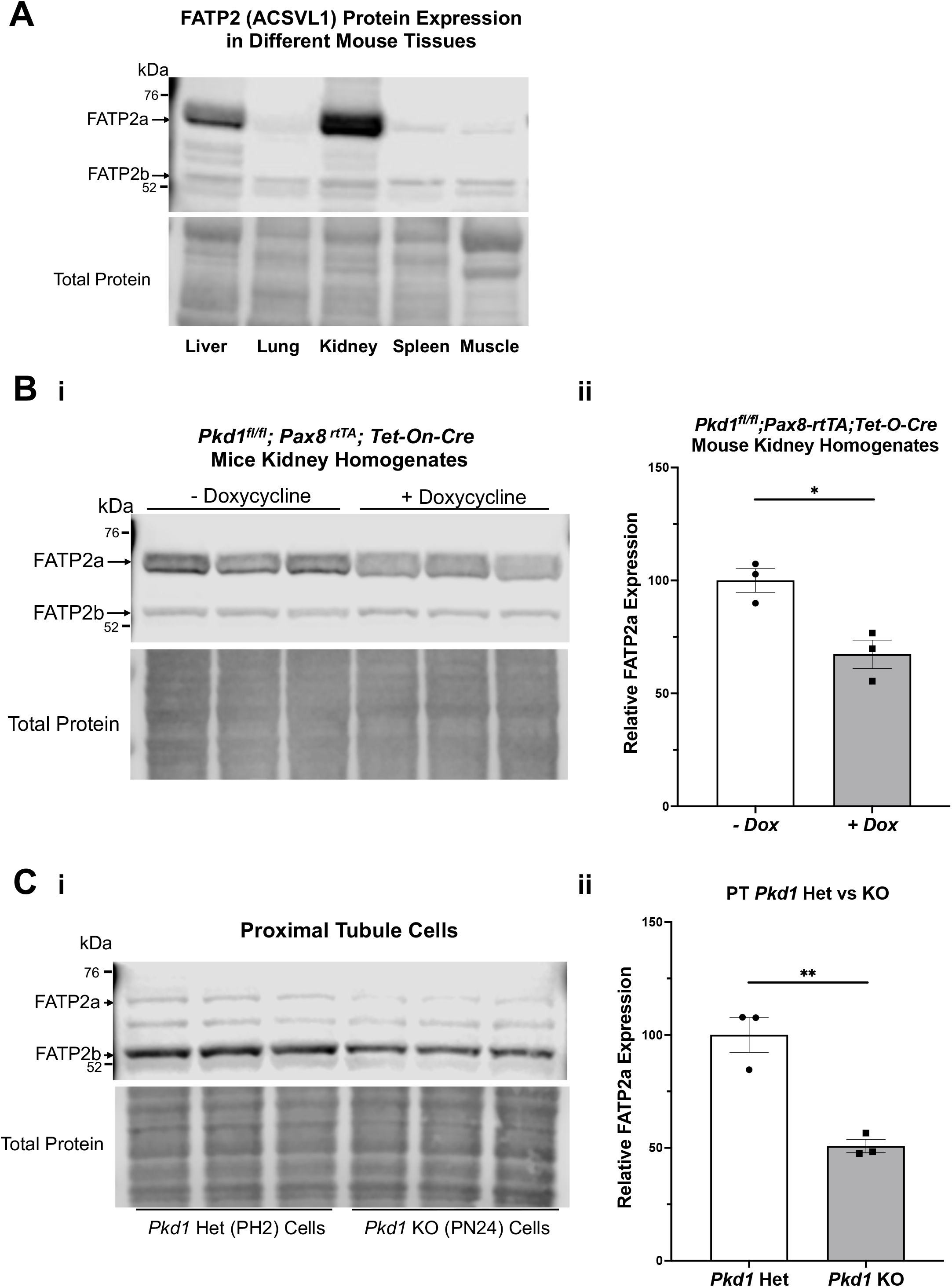
The bempedoic acid-activating enzyme ACSVL1 (FATP2) is expressed in different mouse tissues and PKD kidneys and kidney cell lines. **A**. *Upper*, immunoblotting of various mouse tissues reveals expression of two distinct FATP2 isoforms. The short splice variant (FATP2b), which lacks the acyl-CoA synthetase domain required for conversion of BA to its active metabolite BA-CoA, is expressed in all tissues tested. However, the full-length long form (FATP2a at ∼70kDa) has significant expression only in liver and kidney tissue, thus providing specificity of BA pro-drug conversion to its active form only in these tissues. *Lower*, staining for total protein as a loading control. **B**. FATP2 immunoblotting of representative *Pkd1*^*fl/fl*^*;Pax8-rtTA;Tet-O-Cre* mouse kidneys treated with or without doxycycline to induce tubule-specific *Pkd1* gene inactivation at P10-P11. **i**. *Upper*, the immunoblot revealed expression of FATP2a in all mouse kidneys at the time of euthanasia (P22). *Lower*, staining for total protein as a loading control. **ii**. Densitometric analysis of the FATP2a band normalized to total protein revealed a ∼35% reduction of the expression of the active enzyme in *Pkd1*-null mouse kidneys (\**P* < 0.05). **C**. ^*-*^Both FATP2a and FTAP2b isoforms are also expressed in immortalized mouse kidney epithelial cells that were derived from PT. **i**. *Upper;* We observed generally lower FATP2a expression in *Pkd1* KO cells (*right*) than in controls (*left*). *Lower*, staining for total protein as a loading control. **ii**. Densitometric quantification of the FATP2a levels, normalized to total protein indicates that the *Pkd1*-null PT cells express approximately 50% less FATP2a than the heterozygous *Pkd1*^*+/-*^ cells (\*\**P* < 0.01). Three representative lysate immunoblots are shown for each condition.

### Pkd1 knockout in kidney cell lines is associated with an increase in active ATP-citrate lyase when compared to control parental cell lines

Due to the abundance of ACLY expression in non-cystic PT cells, it was challenging to evaluate differences in ACLY expression and activity by immunoblot of total kidney homogenates. To assess whether cell lines that recapitulate ADPKD cystic disease ex vivo express ACLY, one of the targets of BA, we tested the expression of this enzyme by immunoblotting PT- and IMCD-derived model cell lines in cells grown in 2D. All four cell lines tested, parental PT-derived *Pkd1*^+/-^ (PH2) cells, PT-derived *Pkd1*^*-/-*^ (PN24) cells (Joly et al., 2006) and IMCD-derived WT-IMCD3 and ID1-3E5 *Pkd1*^-/-^ cells (Decuypere et al., 2021) express ACLY (total ACLY or tACLY; **Figure 3A** and **C**). In addition, these cell lines also expressed an active form of ACLY phosphorylated at a target site of Akt (also known as Protein Kinase B; PKB) at Ser-455 (pS455 ACLY). We then normalized the levels of active pS455 ACLY to tACLY in the same samples. We found a significant increase of active ACLY in the *Pkd1*^*-/-*^ cells from both PT and IMCD origin, compared to parental controls (**Figure 3B** and **D**). These results suggest that ACLY activity is increased in ADPKD epithelial cells with cyst-forming capability, compared to cells that are heterozygous for *Pkd1* KO or wild-type at the *Pkd1* locus. Moreover, we surmised that the higher ACLY activity in these ADPKD cell models potentially contributes to cystic growth and disease progression, and thus that targeting this enzyme for inhibition by BA in vivo may be beneficial in reducing cyst size in ADPKD mouse models.

**Figure 3.**
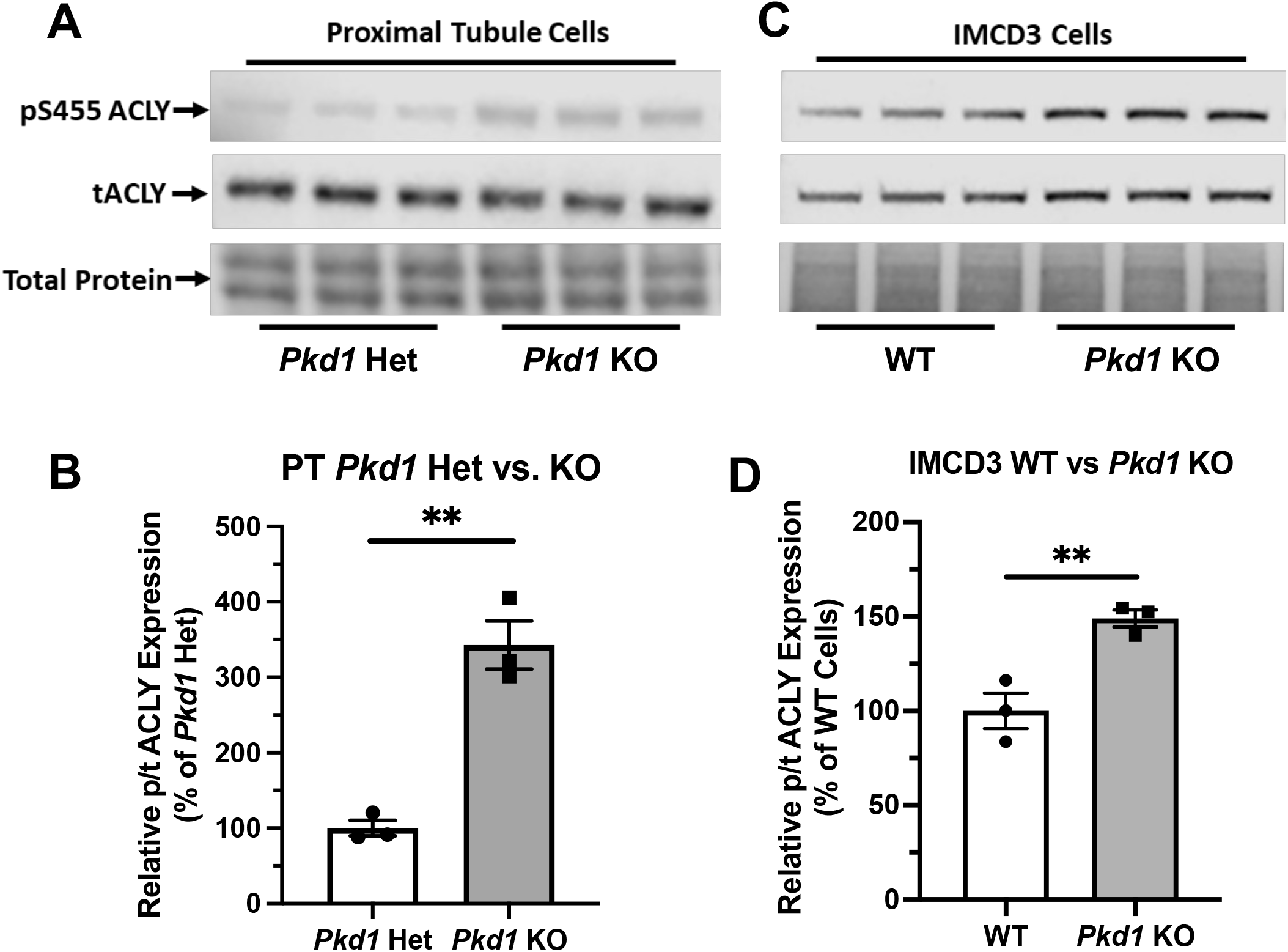
ACLY activity is increased in *Pkd1*^*-/-*^ kidney epithelial cells as compared to controls. **A**. Immunoblotting of proximal tubule (PT)-derived mouse epithelial cells probed for total ACLY expression (tACLY, *upper*) and activated ACLY, as detected using a phospho-specific antibody directed against the Akt phosphorylation site Ser455 (pS455 ACLY, *middle*), in PN24 cells with *Pkd1* expression knocked out at both alleles as compared with PH2 cells heterozygous for *Pkd1* deletion. *Lower*, total protein staining as loading control. **B**. Densitometric quantitation of the pS455 ACLY/tACLY ratio shows a significant ∼250% increase in ACLY activation in the *Pkd1*-null cells as compared to *Pkd1*-het controls (\*\**P*<0.01; unpaired t-test; n = 3). **C**. Immunoblotting of inner medullary collecting duct (IMCD)-derived mouse epithelial cells for expression of tACLY (*upper*) and pS455 ACLY (*middle*), in *Pkd1*-null (ID1-3E5) cells as compared with WT (IMCD3). *Lower*, total protein staining as loading control. **D**. Densitometric quantitation revealed a significant ∼50% increase in the pS455 ACLY/tACLY ratio in the ID1-3E5 cells relative to control IMCD3 cells (\*\**P* < 0.01; unpaired t-test; n = 3).

### Bempedoic acid treatment and the ATP-citrate lyase inhibitor SB-204990 dramatically inhibited cystic growth in Pkd1-null kidney cells lines grown under 3D cyst-forming conditions

We generated cysts from PT-derived *Pkd1*^*+/-*^ (PH2) control cells and *Pkd1*^*-/-*^ (PN24) ADPKD model kidney epithelial cell lines by growing the cells in Matrigel for a total of 12 days and treating these cells with forskolin plus a phosphodiesterase inhibitor (IBMX) from day 2-12. This treatment enhances cyst formation in these commonly used cell culture models (Cabrita et al., 2020). Once cysts appeared by day 9, we exposed the PN24 cultures to either vehicle control (DMSO), the AMPK activator and ACLY inhibitor BA, or the ACLY inhibitor SB-204990 (Chu et al., 2010) for the last three days of the experiment. Under light microscopy, we observed only minimal to very small cystic structures in PH2 control cultures (**Figure 4Ai**, *left*) as compared to PN24 (*Pkd1*^*-/-*^) ADPKD cultures, which resulted in much larger cystic structures (**Figure 4Ai**, *right*). We quantified the cystic area from those micrographs using ImageJ software, and as shown in **Figure 4Aii**, there was a dramatic increase in cyst size in the *Pkd1-*null cells (PN24) as compared to the *Pkd1*^+/-^ heterozygous cells under cyst-inducing conditions. Moreover, in PN24 (*Pkd1*^*-/-*^) ADPKD cultures there was a substantial decrease in cyst size in the cultures treated with BA (**Figure 4Bi**, *middle*) or SB-204990 (**Figure 4Bi**, *right*) as compared with vehicle control (**Figure 4Bi**, *left*). Using the same technique as in **Figure 4Aii**, in ImageJ software, we quantified the cystic area from those micrographs. As summarized in **Figure 4Bii**, there was a significant reduction in cyst area in the BA- and SB-204990-treated cultures as compared with those exposed to vehicle control. We similarly tested whether these two ACLY inhibitors could reduce cyst area in IMCD-derived *Pkd1*^*-/-*^ (ID1-3E5) cells grown in 3D Matrigel cultures treated with IBMX and forskolin and found that BA and SB-204990 both significantly reduced cyst size compared with vehicle (DMSO) (**Supplementary Figure 2**).

**Figure 4.**
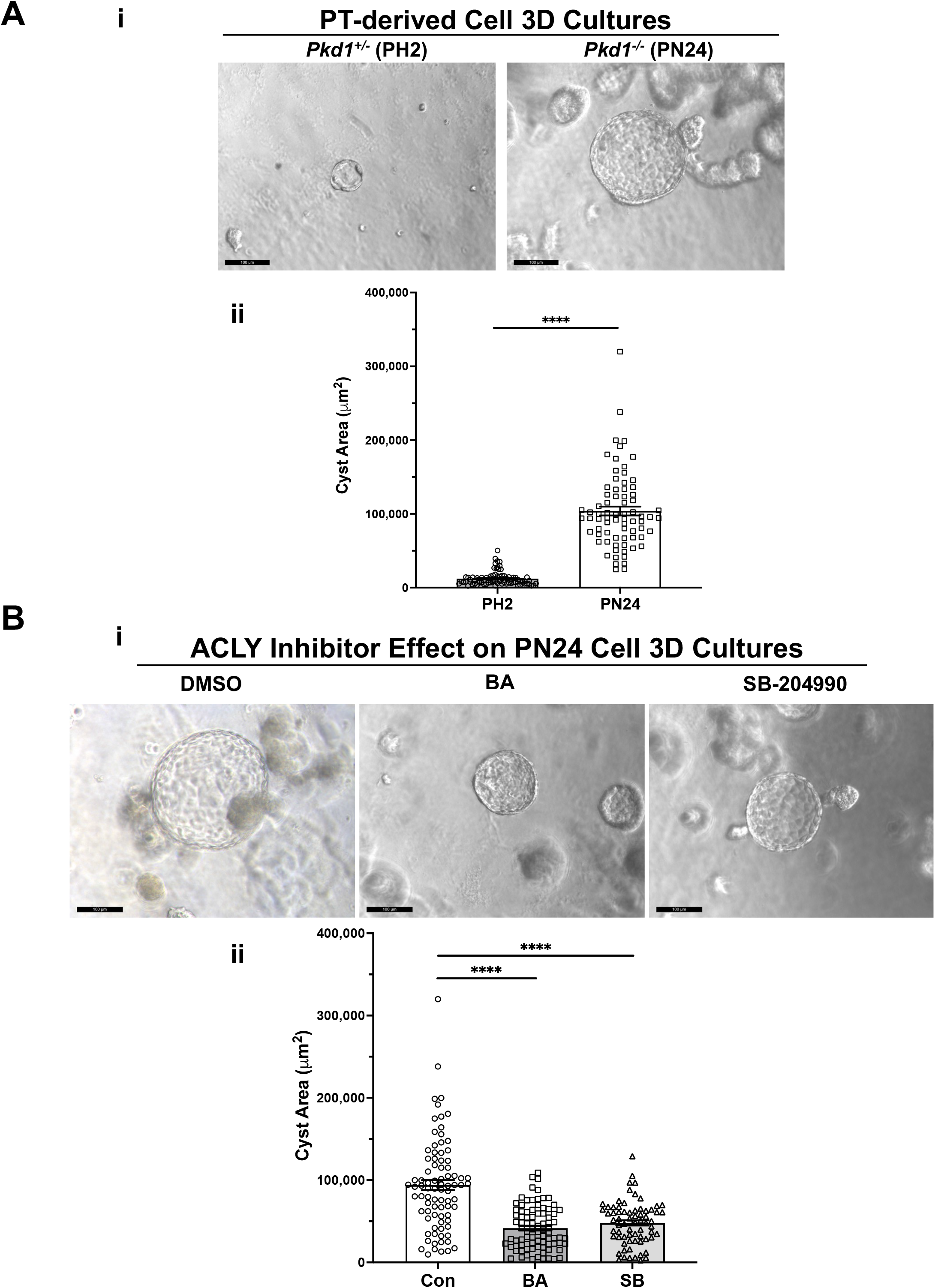
ACLY inhibitors reduce cyst size in 3D cultures of PT-derived *Pkd1*^*-/-*^ (PN24) kidney cells. **A**. PT-derived *Pkd1*^*-/-*^ cells (PN24) and *Pkd1*^*+/-*^ (PH2) cells were grown for 10-12 days in Matrigel supplemented with forskolin + IBMX after 1 day, before imaging and cyst size analysis. **i**. Representative light microscopy images of cystic structures from PH2 vs. PN24 3D cultures (scale bar = 100 µm). **ii**. Summary data reveal that PT-derived *Pkd1*^-/-^ (PN24) cells developed dramatically bigger cystic structures than *Pkd1*^+/-^ (PH2) (n = 74-82, in a total of 4 independent experiments (\*\*\*\**P* < 0.0001). **B**. ACLY inhibitors BA and SB-204990 inhibit cyst growth of PT-derived *Pkd1*^*-/-*^ (PN24) kidney epithelial cells in 3D culture. Cells were cultured for a total of 10-12 days in Matrigel supplemented with forskolin + IBMX after 1 day, and then treated with either vehicle (DMSO; CON), 100 µM BA or 30 µM SB-204990 for the last 3 days of culture before imaging and cyst size analysis. **i**. Representative images of cystic structures in 3D culture of PN24 cells DMSO (*left*) vs. BA (*middle*) vs. SB-204990 treatment (*right*; scale bar = 100 µm). **ii**. Summary data reveal that in PN24 cells treated with BA or SB-204990 the cyst area relative to CON is dramatically reduced (n = 69-84, represent 3-5 independent experiments; \*\*\*\**P* < 0.0001 for the indicated comparisons).

**Figure 5.**
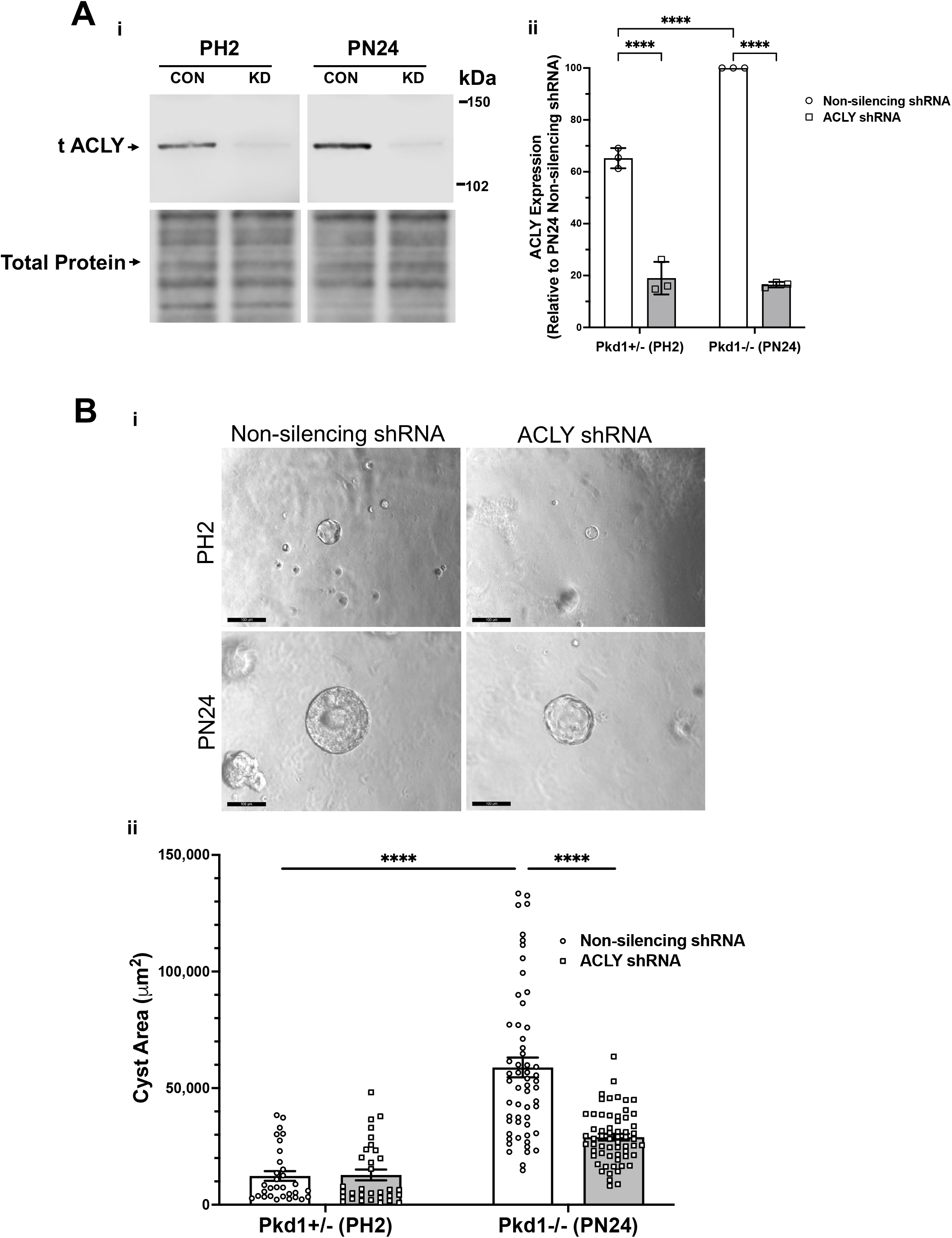
ACLY knockdown inhibits cystic growth in 3D cultures of PT-derived kidney epithelial cells. **A.** Stable PT-derived cell lines expressing either non-silencing control shRNA or shRNA against ACLY were generated and checked for ACLY protein expression by immunoblotting analysis. i. Representative immunoblotting of ACLY protein expression in the different cell lines. ii. Summary data reveal that *Pkd1*^*+/-*^ (PH2) cells expressing non-silencing control shRNA have ∼35% reduced ACLY expression compared with ACLY expression in *Pkd1*^*-/-*^ (PN24) cells expressing non-silencing control shRNA. There were means of 71% and 83% knockdown of ACLY expression in PH2 and PN24 cells, respectively, expressing shRNA against ACLY relative to ACLY expression in cells expressing non-silencing control shRNA (n = 3, \*\*\*\**P* < 0.0001 for the indicated comparisons). **B**. *Pkd1*^*-/-*^ (PN24) cells developed significantly larger cysts than *Pkd1*^*+/-*^ (PH2) cells, and shRNA-mediated ACLY knockdown inhibited cyst growth of *Pkd1*^*-/-*^ (PN24) kidney epithelial cells in 3D culture. Cells were cultured for a total of 12 days in Matrigel supplemented with forskolin + IBMX after day 1, and cysts were imaged and cyst size was analyzed as described in *Materials and Methods*. i. Representative images of cystic structures in the different cell lines. ii. Summary data reveal *Pkd1*^*-/-*^ (PN24) cells developed significantly larger cysts than *Pkd1*^*+/-*^ (PH2) cells (n = 31-57 cysts analyzed from 3 biological replicate experiments; \*\*\*\**P* < 0.0001). Cells expressing shRNA directed against ACLY dramatically reduced mean cystic areas relative to those of cells expressing non-silencing control shRNA (n = 57-61 cysts analyzed from 3 biological replicate experiments, \*\*\*\**P* < 0.0001).

**Figure 6.**
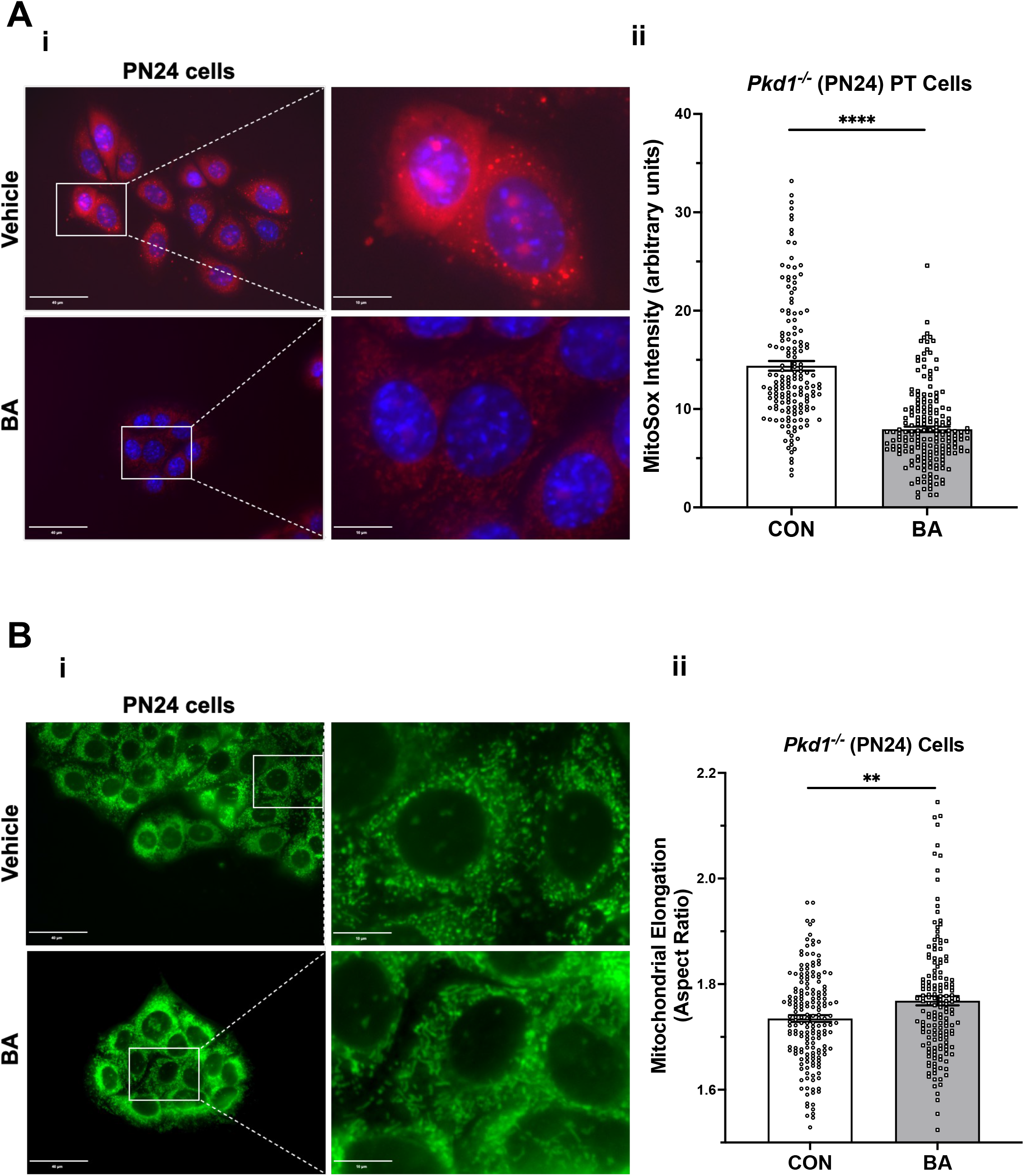
Bempedoic acid inhibits mitochondrial superoxide production and promotes mitochondrial elongation in *Pkd1*^*-/-*^ cells. **A**. To analyze the effect of BA on mitochondrial superoxide production, PT-derived *Pkd1*^*-/-*^ (PN24) cells were stained with MitoSOX™ Red mitochondrial superoxide indicator (*red*) and Hoechst 33342 nuclear stain (*blue*). **i**. Representative epifluorescence micrograph images are shown of PN24 cells in the absence (*top*) or presence (*bottom*) of BA treatment (100 µM) for 24 h. Left panel scale bars = 40 µm. Right panels show enlargement of inset areas in the left panels (right panel scale bars = 10 µm). **ii**. Summary data reveal that BA treatment dramatically decreased mitochondrial superoxide production in PN24 cells (n = 159-188 cells analyzed from four biological replicates; \*\*\*\**P*<0.0001). **B**. BA treatment significantly increased mitochondrial elongation of *Pkd1*^*-/-*^ PN24 cells. **i**. Representative images of MitoTracker Green-stained cells in the absence (*top*) or presence (*bottom*) of BA treatment (100 µM) for 24 h. Left panel scale bars = 40 µm. Right panels show enlargement of inset areas in the left panels (right panel scale bars = 10 µm). **ii**. Summary data reveal that BA treatment significantly increased mean cellular mitochondrial elongation in PN24 cells (mean cellular mitochondrial elongation values on n = 159-177 cells analyzed from four biological replicates, as described in *Materials and Methods*; \*\**P*<0.01).

### Acly knockdown inhibited cystic growth in Pkd1-null kidney cell 3D cultures

To examine the role of ACLY more directly in governing ADPKD cyst growth in vitro, we generated stably transduced PH2 and PN24 cell lines using the pGIPZ lentiviral system to express either shRNA directed against mouse *Acly* (KD cells) or a non-silencing shRNA control (**Figure 5**). ACLY protein expression in both the PN24 (*Pkd1*^*-/-*^) and PH2 (*Pkd1*^*+/-*^) cells was dramatically inhibited to ∼20-30% of levels in the corresponding non-silencing controls, and baseline ACLY expression was significantly reduced in the PH2 cells relative to PN24 cells (**Figure 5A**). We then performed 3D cyst growth assays in these cells as described above. While PH2 cells generated only small cystic structures whose size was not significantly affected by *Acly* KD, cyst growth in the PN24 *Acly* KD cells was significantly inhibited relative to that in the PN24 non-silencing control cells (**Figure 5B**). Taken altogether, these experiments further confirmed our hypothesis that BA treatment and ACLY inhibition has the promise of a therapeutic effect in vivo by inducing a significant reduction in cyst size in ADPKD in vitro 3D cell culture models.

### Bempedoic acid inhibits mitochondrial superoxide production and promotes mitochondrial elongation in ADPKD kidney-derived cell lines

Normal mitochondrial function is regulated in part by the interplay of two targets of BA, AMPK and ACLY (Cf. **Figure 1**). These key metabolic enzymes appear to be dysregulated in ADPKD, where there is upregulation of ACLY activity (**Figure 3**) and downregulation of AMPK activity (Rowe et al., 2013, Song et al., 2020). Moreover, when mitochondria are unable to efficiently utilize fatty acids for oxidative metabolism (Cf. **Figure 1 - step 6**) with decreased electron transfer in the respiratory chain, there is increased reactive oxide species (ROS) production leading to accumulation of superoxide (Lenaz, 2001). Mitochondrial dysfunction with increased oxidative stress is a key feature of ADPKD (Chang and Ong, 2018).

Moreover, it has been shown that AMPK activation may improve mitochondrial function in ADPKD and overall disease severity (Song et al., 2020). BA would thus appear to be a good candidate drug to induce simultaneous ACLY inhibition and AMPK activation as regulators of mitochondrial function.

To examine whether BA-mediated cyst reduction is associated with improved mitochondrial function in ADPKD, we first tested the effects of BA on mitochondrial superoxide production in PT-derived *Pkd1*^*-/-*^ (PN24) cells (**Figure 6A**). These cells were loaded with MitoSOX™ Red, a mitochondrial superoxide indicator, and incubated in the presence of BA or vehicle for 24 h. Our results support the hypothesis that BA improves mitochondrial function, as demonstrated by reduced fluorescence signal from MitoSOX™ Red in cells treated with BA compared to the vehicle control. We found a similar reduction in MitoSOX™ Red intensity in IMCD-derived *Pkd1*^*-/-*^ (ID1-3E5) cells treated with BA relative to vehicle control (see **Supplementary Fig. 3A**). These findings indicate that there is reduced mitochondrial superoxide production with BA treatment relative to controls in both PT-derived *Pkd1*^*-/-*^ (PN24) or IMCD-derived *Pkd1*^*-/-*^ (ID1-3E5) cells.

Mitochondrial morphology is another important indicator of mitochondrial oxidative function. Specifically, mitochondrial elongation facilitates cristae formation and assembly of respiratory complexes to enhance oxidative phosphorylation in cells (Li et al., 2017). Germino and colleagues demonstrated that mitochondrial elongation is defective in ADPKD model cell lines with decreased elongation in *Pkd1*-null cells compared to wild-type controls (Lin et al., 2018), as reviewed in (Padovano et al., 2018). Here we tested the effects of BA treatment on mitochondrial elongation (or aspect ratio) using MitoTracker™ staining in both PT-derived PN24 cells (**Figure 6B**) and IMCD-derived ID1-3E5 cells (**Supplementary Fig. 3B**). Quantification of mitochondrial length divided by width was performed on the IF images using Image J. We found that BA treatment significantly increased mitochondrial elongation of both *Pkd1*-null cell lines (**Figure 6Bii** and **Supplementary Fig. 3Bii**). These findings are consistent with the hypothesis that BA promotes mitochondrial oxidative function in *Pkd1*-deficient cells.

### In vivo studies testing the effects of bempedoic acid and tolvaptan on PKD disease severity in an early/rapid induced Pkd1 gene inactivation ADPKD mouse model

The use of tolvaptan, the only FDA-approved drug currently to treat ADPKD progression in patients with certain characteristics, is limited by its side effects, such as polyuria and thirst, potential hepatotoxicity, and its availability and cost (Blair, 2019). Future ADPKD treatment strategies may involve combination therapies including tolvaptan and other drugs that target complementary dysregulated cellular signaling pathways, potentially conferring synergistic or additive benefits, and allowing lower drug doses than those used in ADPKD monotherapy. To evaluate the effects of BA and tolvaptan in vivo we used an early and rapid model of ADPKD progression, the *Pkd1*^*fl/fl*^; *Pax8-rtTA; Tet-O-Cre* mouse. The disease was induced via doxycycline administration at P10-P11 to inactivate *Pkd1*. Subsequently, the mice were treated with either vehicle, BA, BA plus tolvaptan or tolvaptan alone through P21 prior to euthanasia at P22, a timeline summarized in **Figure 7A**. These mice progress to ESKD at around the time of euthanasia (Ma et al., 2013). We first measured changes in the % total kidney weight/body weight (%TKW/BW) and blood urea nitrogen (BUN) levels at the time of euthanasia as markers of disease severity in this ADPKD model. Representative kidney section micrographs stained by H&E under the different treatment conditions are shown in **Figure 7B**. As shown in **Figure 7C**, there were graded reductions of the %TKW/BW that occurred with the BA and/or tolvaptan treatments. Specifically, BA (30 mg/kg/d) reduced %TKW/BW vs. vehicle at euthanasia (6.9 vs. 11.9%; *P*<0.01). Similarly, tolvaptan (100 mg/kg/d) reduced %TKW/BW to 7.8% vs. vehicle (*P*<0.05). Addition of BA (30 mg/kg/d) to tolvaptan (100 mg/kg/d) caused a further reduction in %TKW/BW (4.9%; *P*<0.05) vs. tolvaptan alone. As shown in **Figure 7D**, there were also graded reductions of BUN levels at euthanasia with BA and/or tolvaptan treatments. BA treatment was associated with a reduced levels of BUN vs. vehicle (59 vs. 107 mg/dL; *P*<0.05). Tolvaptan treatment is associated with lower levels of BUN relative to vehicle at 30 (68 mg/dL; *P*<0.05) and 100 mg/kg/d (35 mg/dL; *P*<0.01). Again, addition of BA to tolvaptan at 30 mg/kg/d led to further significant reduction in BUN (38 mg/dL; *P*<0.05). In summary, these findings indicate that both BA and tolvaptan reduce kidney growth and improve kidney function in this early onset ADPKD mouse model, and there are additive benefits with combination therapy.

**Figure 7.**
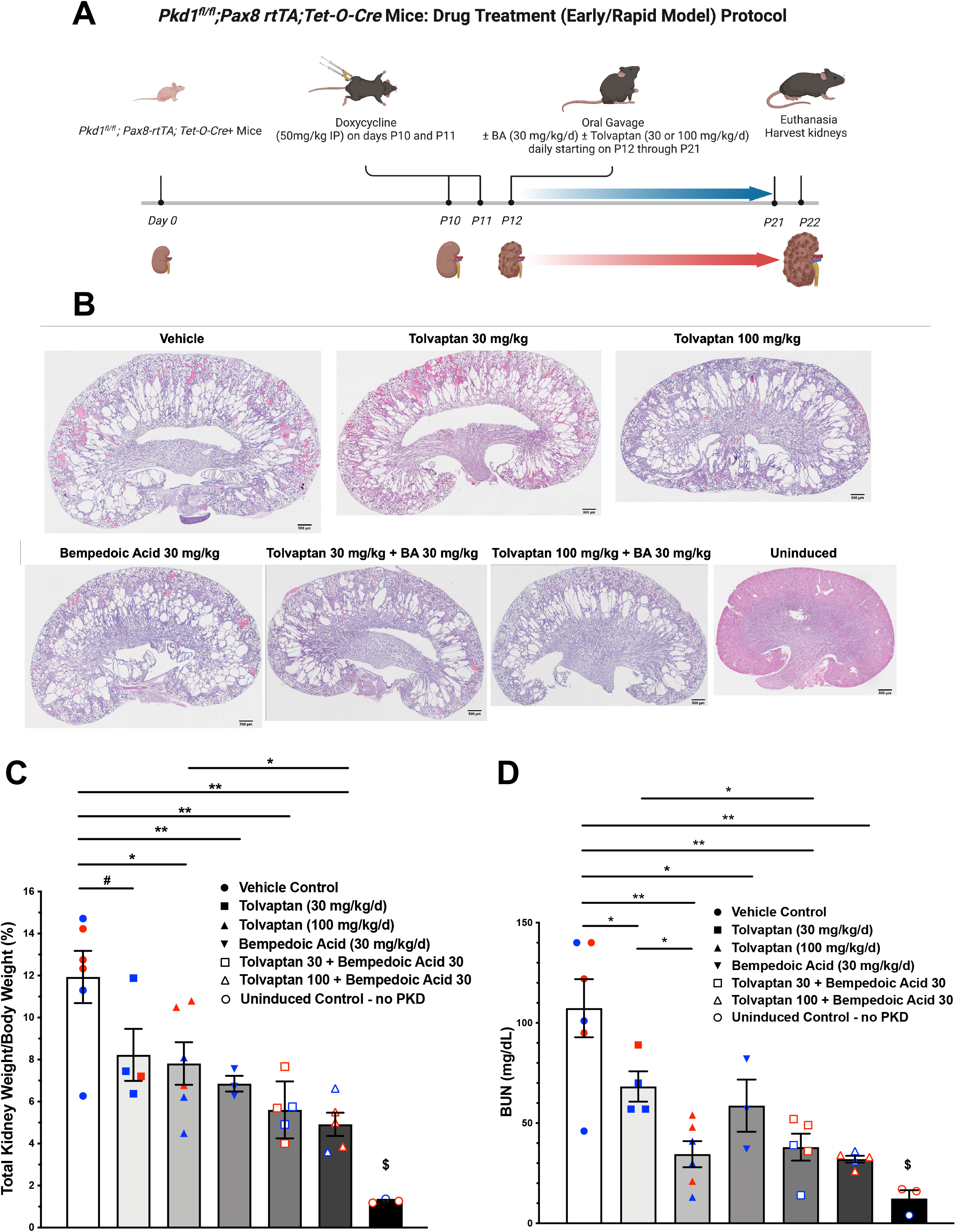
Bempedoic acid treatment alone and in combination with tolvaptan reduces PKD disease severity in an early/rapid induced *Pkd1* gene inactivation ADPKD mouse model. As measures of disease severity in vivo, *Pkd1*^*fl/fl*^; *Pax8-rtTA; Tet-O-Cre* mice were induced with doxycycline to inactivate *Pkd1* and then treated ± BA ± tolvaptan for 10 days prior to euthanasia, as shown schematically (**A**), and then total kidney weight/body weight ratio (TKW/BW) and blood urea nitrogen (BUN) levels by iStat were evaluated as described in *Materials and Methods*. **B**. Representative H&E-stained sagittal kidney sections under the different treatment conditions are shown. BA (30 mg/kg/d) reduced mean (± SE) TKW/BW (**C**) and BUN (**D**) to a similar extent as tolvaptan (30 and 100 mg/kg/d). Addition of BA to tolvaptan caused further reductions in TKW/BW and BUN vs. tolvaptan alone (\**P* < 0.05, \*\**P* < 0.01, and ^#^0.05 < *P* < 0.10 for the indicated comparisons). ^$^Significantly different from all other treatment conditions. Data obtained from male and female mice were combined for each treatment condition as mice were studied at ages before reaching reproductive capability. Blue and red data points shown in panels C and D correspond to male and female mice, respectively.

### Effects of bempedoic acid and tolvaptan treatment in PKD mice on protein expression of key signaling and injury markers in kidney homogenates

Kidneys from the conditional *Pkd1* knockout mice were harvested at the time of euthanasia to analyze the effects of treatment with BA and tolvaptan on relevant target proteins (**Figure 8**). As expected, BA treatment generally reduced ACLY activity (pSer455 ACLY (pACLY); **Figure 8A**) and stimulated AMPK activity (pThr172 AMPK (pAMPK); **Figure 8B**) in kidney tissue homogenates relative to vehicle controls or treatment with tolvaptan alone. BA also tended to inhibit mTOR and ERK pathway signaling, which are upregulated in ADPKD (Saigusa and Bell, 2015), as evidenced by decreased phosphorylation of P70S6K (**Figure 8C**) and ERK (**Figure 8D**), respectively, relative to paired controls in the absence of BA. Finally, BA also sharply reduced expression of the PT kidney injury marker KIM1 (**Figure 8E**) and, to a lesser extent, tended to inhibit the distal kidney injury marker NGAL (**Figure 8F**) relative to controls. These BA effects occurred both alone and in combination with tolvaptan.

**Figure 8.**
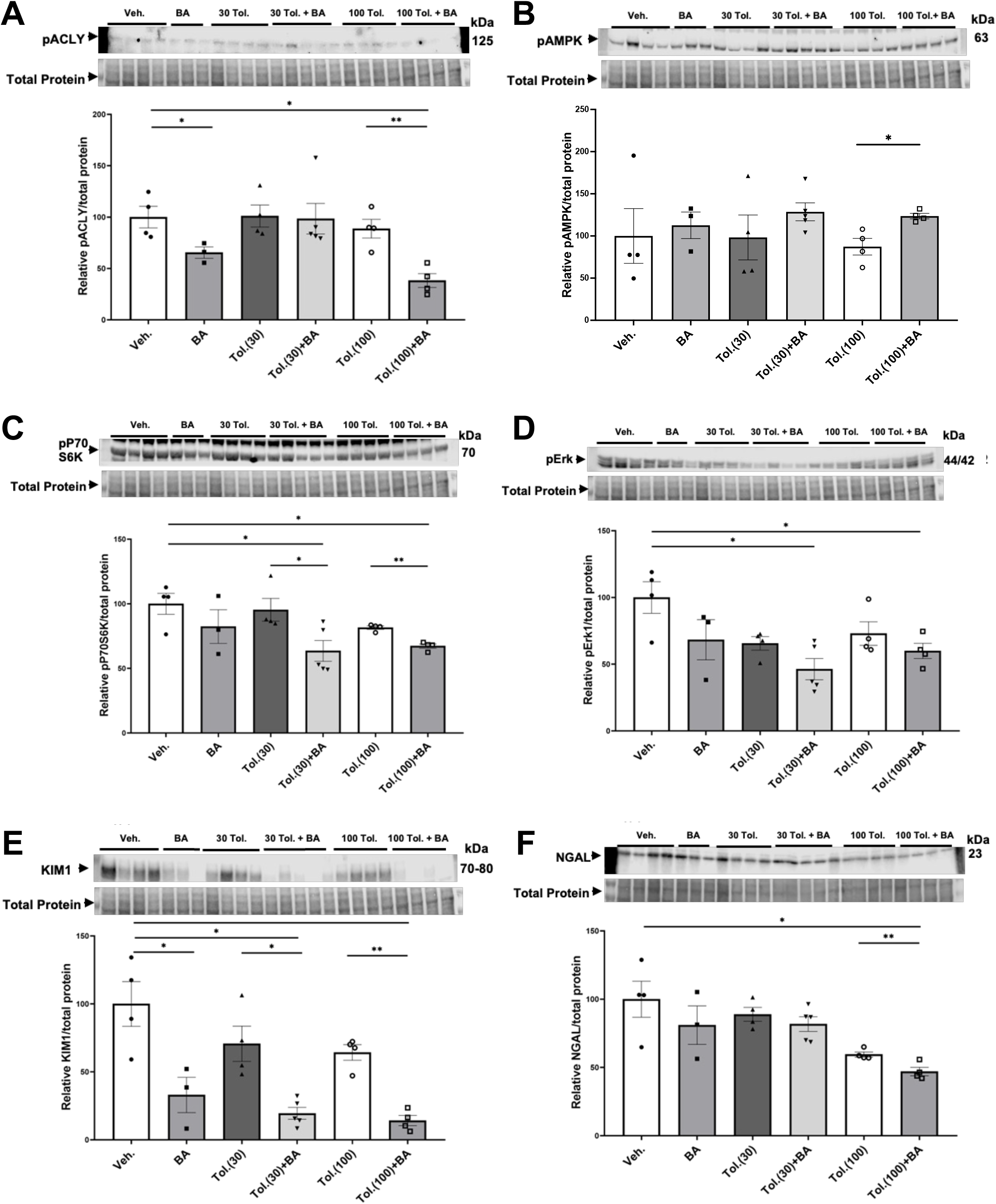
Effects of bempedoic acid and tolvaptan treatment in PKD mice on protein expression of key signaling and injury markers in kidney homogenates. Immunoblots were performed on kidney tissue homogenates from *Pkd1*^*fl/fl*^; *Pax8-rtTA; Tet-O-Cre* mice with early/rapid induced *Pkd1* gene inactivation with or without concurrent treatment with BA (30 mg/kg/d) and/or tolvaptan (30 or 100 mg/kg/d). Densitometric intensities of immunoblot bands from each lane were normalized to the total protein signal from that lane as a loading control (shown in lower panels of each immunoblot). Kidney homogenates were probed for ACLY activity (pSer455 ACLY (pACLY); **A**), AMPK activity (pThr172 AMPK (pAMPK); **B**), mTOR activity (pP70S6K; **C**), ERK activity (pERK; **D**), and the kidney injury markers, KIM1 (**E**), and NGAL (**F**) (\**P* < 0.05, \*\**P* < 0.01 for the indicated comparisons). Representative immunoblots are shown at top and summary quantitation of mean (± SE) relative protein expression levels are shown at bottom of each panel.

### Effects of bempedoic acid and tolvaptan treatment in ADPKD mice on protein expression of key signaling markers in liver homogenates

The liver is a main target for BA in its inhibition of sterol synthesis and its other metabolic effects (Pinkosky et al., 2016). As idiosyncratic hepatotoxicity is a concern in ADPKD patients treated with tolvaptan (Blair, 2019), we also examined protein expression of several markers in liver homogenates from the conditional *Pkd1* knockout mice to analyze the effects of treatment with vehicle control, BA (30 mg/kg/d) in the presence or absence of high-dose tolvaptan (100 mg/kg/d; **Figure 9**). BA treatment tended to cause the expected decrease in ACLY activity (pACLY) in liver tissue (**Figure 9A**) while increasing AMPK activity, as evidenced by increased pAMPK (**Figure 9B**). BA also tended to increase phosphorylation of acetyl-CoA carboxylase (pACC), a canonical target of AMPK (**Figure 9C**). Interestingly, BA treatment generally caused an enhancement in the expression in liver of FATP2, the fatty acid transporter that is required for activation of the BA pro-drug (Pinkosky et al., 2016), both alone and in combination with tolvaptan (**Figure 9D**). This effect of BA treatment enhancing FATP2 expression was also found in kidney tissue homogenates (**Supplementary Fig. 4**). Of note, BA also appears to promote mitochondrial biogenesis as measured by enhanced expression of PGC-1α in the ADPKD livers, while high-dose tolvaptan tended to decrease PGC-1α (**Figure 9E**). Finally, BA treatment dramatically decreased cleaved caspase-3, a key marker for apoptosis, in ADPKD mice livers, both alone and in combination with tolvaptan (**Figure 9F**). This finding suggests that BA may serve a protective role in preventing apoptosis, a marker of hepatocyte injury and death, in the setting of high-dose tolvaptan treatment.

**Figure 9.**
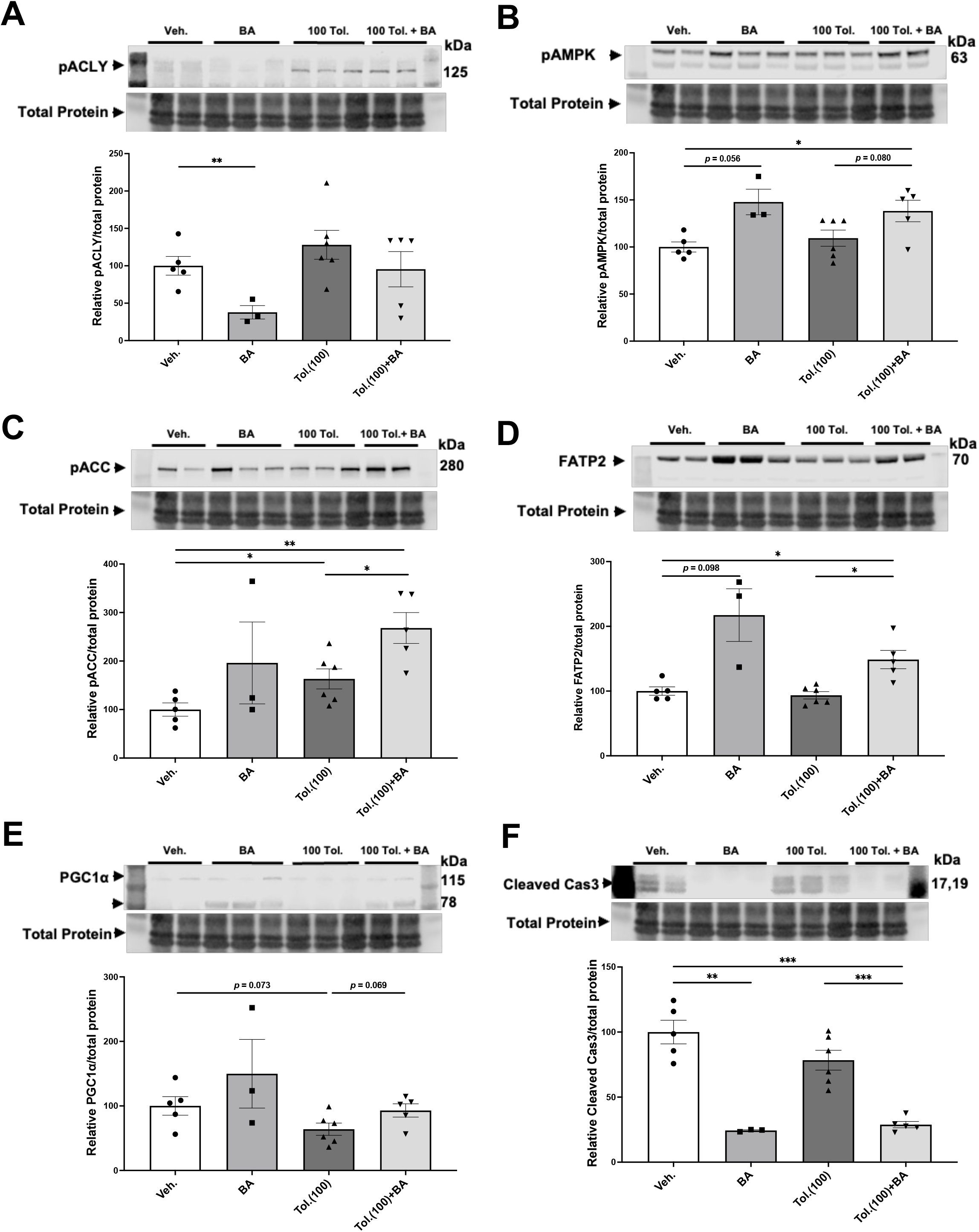
Effects of bempedoic acid and tolvaptan treatment in PKD mice on protein expression of key signaling markers in liver homogenates. Immunoblots were performed on liver tissue homogenates from *Pkd1*^*fl/fl*^; *Pax8-rtTA; Tet-O-Cre* mice with early/rapid induced *Pkd1* gene inactivation with or without concurrent treatment with BA (30 mg/kg/d) and/or tolvaptan (100 mg/kg/d). Densitometric intensities of immunoblot bands from each lane were normalized to the total protein signal from that lane as a loading control. Liver homogenates were probed for ACLY activity (pSer455 ACLY; **A**), AMPK activity (pThr172 AMPK (pAMPK; **B**) and pACC (**C**)), FATP2 expression (**D**), the mitochondrial biogenesis marker PGC-1α (**E**), and the apoptosis marker cleaved caspase 3 (**F**) (\**P* < 0.05, \*\**P* < 0.01, \*\*\**P* < 0.001 or as stated for the indicated comparisons). Representative immunoblots are shown at top and summary quantitation of mean (± SE) relative protein expression levels are shown at bottom of each panel.

## DISCUSSION

There are few therapeutic options for people living with ADPKD to arrest disease progression. As it has become recognized that ADPKD is a metabolic disease with dysregulated mitochondrial function (Padovano et al., 2018), identifying new therapies that target dysregulated metabolism are especially attractive. In searching for new metabolic targets to ameliorate the disease and prevent its progression, we found that ACLY activity was increased in PT- and IMCD-derived *Pkd1* KO cells relative to controls (**Figure 3**). As BA was recently FDA-approved for treatment of hypercholesterolemia, we considered that repurposing this drug could be compelling to test as a novel ADPKD therapeutic. Consistent with previous work (Pinkosky et al., 2016, Steinberg et al., 1999), we confirmed that mouse kidney and liver tissue, along with ADPKD cells, express the FATP2 (ACSVL1) enzyme that mediates the activation of BA (**Figure 2**). Thus, BA should only be converted to its active form in kidney and liver, the two organs primarily affected with cysts in ADPKD. This targeting specificity would be advantageous in ADPKD to reduce potential side effects in other organs, as ADPKD patients may need to be treated for several decades. Indeed, off-target side effects have historically limited the use of certain promising experimental drugs that would appear to have therapeutic benefits in ADPKD (e.g., rapamycin and analogues) (Serra et al., 2010).

For this initial study to explore the potential utility and efficacy of BA as a new therapy for ADPKD, we opted to test its effects first in PKD cell lines and in an inducible ADPKD mouse model that develops early, severe kidney disease. We found that both BA and a distinct ACLY inhibitor SB-204990 inhibited 3D cystic growth in PT- and IMCD-derived *Pkd1*-null epithelial cells (**Figure 4** and **Supplementary Figure 2**). Moreover, ACLY knockdown similarly inhibited 3D cystic growth in PT-derived *Pkd1*-null epithelial cells (**Figure 5**). In *Pkd1*-null cell lines, BA treatment also reduced mitochondrial superoxide production, a marker of cellular oxidative stress, and increased mitochondrial elongation, a marker of mitochondrial oxidative function (**Figure 6** and **Supplementary Figure 3**). To test whether BA is effective in ADPKD to reduce cyst size in vivo, we used an early *Pkd1* conditional KO mouse model of rapidly progressive kidney cystic disease, similar to that described previously (Ma et al., 2013). In this mouse model, we found that BA treatment reduced kidney size and function (BUN) to a similar extent as tolvaptan relative to untreated controls, and addition of BA to tolvaptan caused a further reduction in both markers of cystic disease severity and kidney function when compared with tolvaptan alone (**Figure 7**). The apparent additive benefits of combination BA plus tolvaptan therapy could have important clinical implications if confirmed in human clinical ADPKD trials as it may afford lower dosing to achieve efficacy and with fewer side effects.

Treatment with BA also inhibited key cellular signaling pathways associated with ADPKD cellular proliferation (mTOR and ERK) (Saigusa and Bell, 2015) and sharply reduced the kidney injury marker KIM1 and, to a lesser extent, NGAL (**Figure 8**). Finally, BA therapy in combination with tolvaptan in the *Pkd1* conditional KO mouse model dramatically inhibited apoptosis (cleaved caspase-3) and tended to increase mitochondrial biogenesis (PGC-1α) in liver tissue relative to tolvaptan treatment alone (**Figure 9**), which suggests that BA combination therapy with higher dose tolvaptan could help mitigate tolvaptan-associated hepatotoxicity. Taken altogether, the findings of this study suggest that BA may be considered a promising new therapeutic candidate for ADPKD, which deserves additional exploration.

Interestingly, BA is a pro-drug that requires activation by the very long-chain acyl-CoA synthetase, a fatty acid transporter also known as ACSVL1 or FATP2, which adds a CoA moiety to free fatty acids in kidney and liver cells (Pinkosky et al., 2016). Our initial intent for FATP2 immunoblotting was to ascertain whether the BA-activating enzyme FATP2a was expressed in cells with *Pkd1* knockout, and therefore whether the pro-drug BA could be useful for in vitro cystogenesis assays. Interestingly, we found that there was a reduction of FATP2a protein levels in both *Pkd1*-null cells and kidney tissue relative to controls (**Figure 2** and **Suppl. Figure 1**). Moreover, treatment of *Pkd1*-null cells with BA increased the levels of FATP2a expression compared to control cells treated with vehicle (**Figure 9D** and **Suppl. Figure 4**). Although in-depth characterization of the mechanism for FATP2 regulation by BA is beyond the scope of our current study, we speculate that expression of FATP2a, as a fatty acid transporter, may be determined by differences in the levels or activities of long-chain fatty acids in the *Pkd1*-null cells as compared to controls. Such differences may reflect the profound metabolic changes induced by *Pkd1* knockout, which are perhaps reversed by treatments that ameliorate cystic disease.

Further studies will be important to explore the potential benefits of BA in more relevant mouse and/or other animal models of ADPKD (e.g., slower onset disease models, including the hypomorphic *Pkd1*^*RC/RC*^ mouse model (Hopp et al., 2012), and in other inducible and *Pkd2* mutant animal models). Employing animal models with slower disease trajectories that encompass the animals’ reproductive ages will also allow exploration of potential sex differences with respect to disease parameters and BA treatment effects in the setting of ADPKD. In the ADPKD model presented here, mice develop early disease and are treated before having reproductive capability, with euthanasia occurring at P22, so sex differences were likely not relevant. It is also currently unclear what are the specific downstream targets or relevant ADPKD cellular signaling pathways of BA. Administration of this drug to mice with more chronic cystic disease will allow a more careful and comprehensive exploration of its effects on cellular pathways and its role in regulating disease progression.

Of note, as BA is already FDA-approved and is generally well-tolerated even in patients that lack hypercholesterolemia (Banach et al., 2020, Powell and Piszczatoski, 2021), initiating human clinical trials for ADPKD may become feasible relatively soon. Importantly, as BA targets different cellular signaling pathways than tolvaptan and other emerging therapies, it will be important to test for benefits of BA both alone and in combination with tolvaptan and other drugs that target complementary cellular signaling pathways in future clinical trials.

## ACKNOWLEDGMENTS

We thank Drs. Michael Caplan and Stefan Somlo at Yale University for the generous provision of the *Pkd1*-null cell lines. We also thank Drs. Terry Watnick and Patricia Outeda at the Maryland PKD Core for sharing of the conditional *Pkd1*-null mice.

## AUTHOR CONTRIBUTIONS

K.R.H., H.L., B.S., E.J.C., S.L.P., and N.M.P-S. conceived and designed research. H.L., B.S., S.S., P.H., J.P., J.W., and V.M. performed experiments. K.R.H., H.L., B.S., S.S., P.H., J.P., V.M. and N.M.P.-S. analyzed data. K.R.H., H.L., B.S., S.S., P.H., J.P., V.M., S.L.P., and N.M.P.-S. interpreted results of experiments. K.R.H., H.L., B.S., and N.P.S. prepared figures. K.R.H. H.L., and N.M.P.-S. drafted manuscript. K.R.H., H.L., B.S., S.L.P. and N.M.P.-S. edited and revised the manuscript.

## SUPPLEMENTARY FIGURE LEGENDS

**Supplementary Figure 1.**
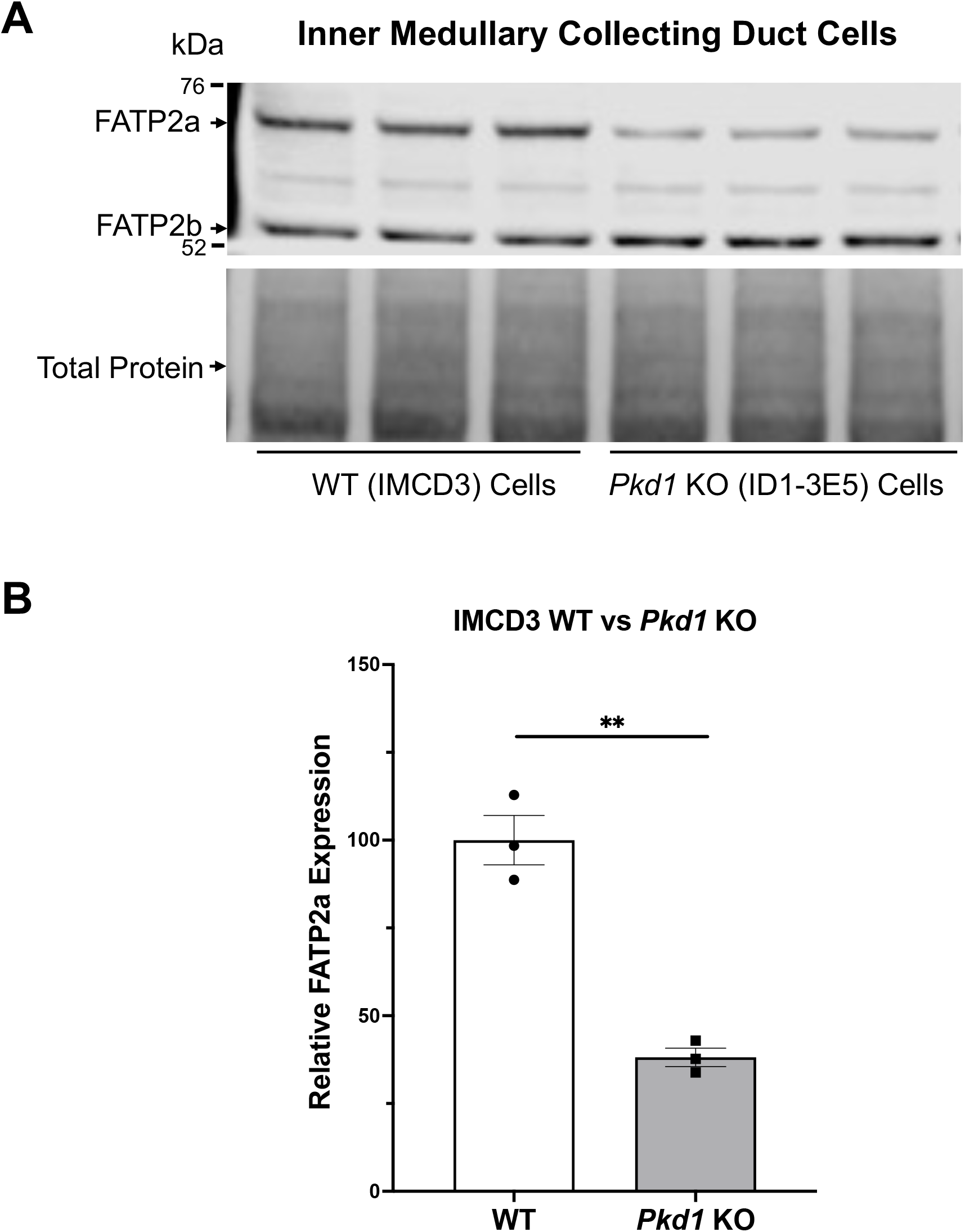
The bempedoic acid-activating enzyme ACSVL1 (FATP2) is expressed in IMCD-derived *Pkd1*^-/-^ kidney cells. **A**. ^*-*^Both FATP2a and FTAP2b isoforms are expressed in this immortalized mouse kidney epithelial cell line of collecting duct origin by immunoblot. FATP2a expression in *Pkd1* KO cells (ID1-3E5, *Upper, right*) than in controls (WT IMCD3, *Upper, left*). *Lower*, total protein signal as loading control for the immunoblot. **B**. Densitometric quantification of the FATP2a levels normalized to total protein indicates that the *Pkd1*-null IMCD-derived cell line (*Pkd1* KO) express approximately 40% less FATP2a than the wild-type IMCD3 cells (WT). Three representative lysate immunoblots are shown for each condition (\*\**P* < 0.01).

**Supplementary Figure 2.**
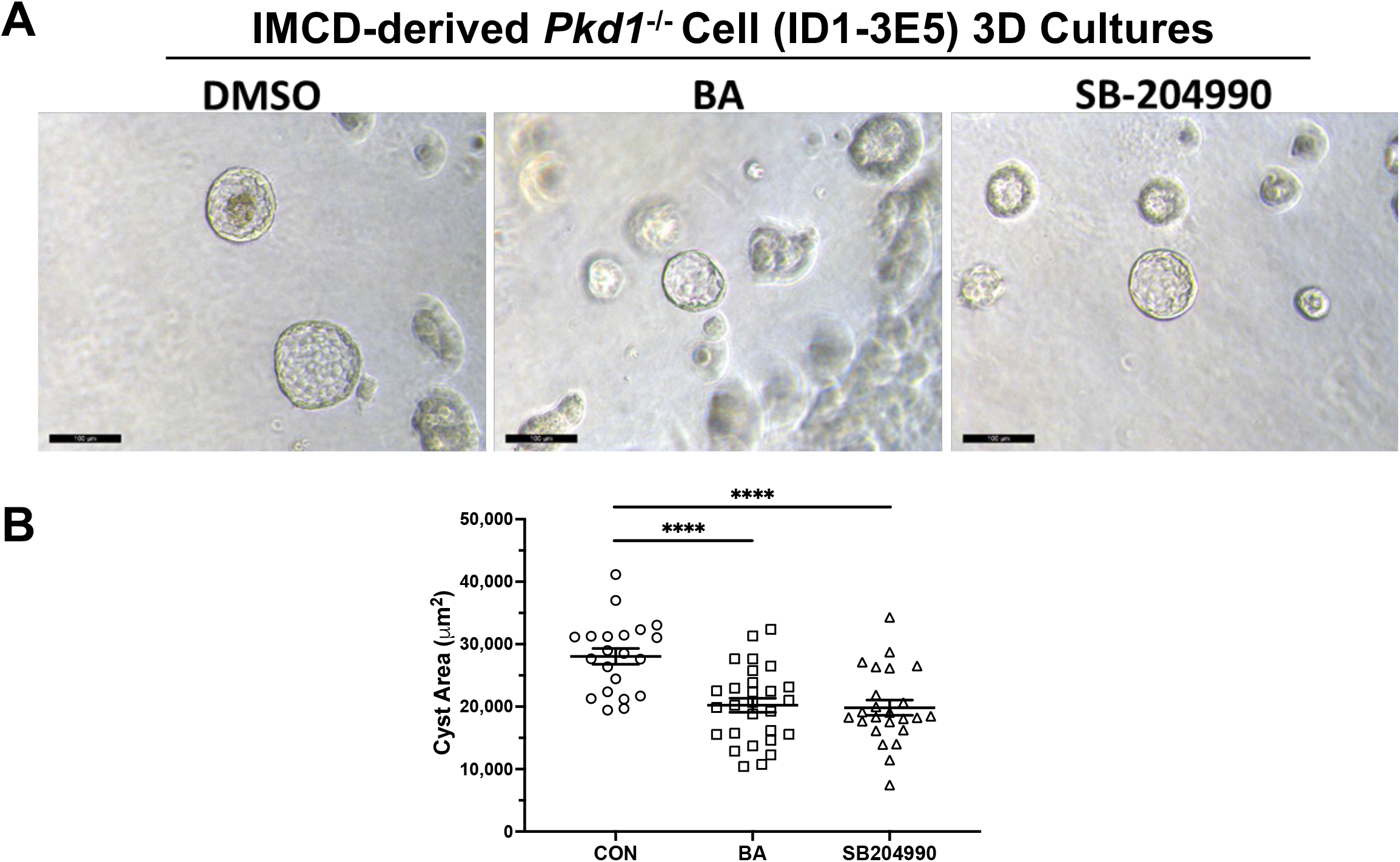
ACLY inhibitors dramatically inhibit cystic growth in 3D cultures in IMCD-derived *Pkd1*^-/-^ kidney cells. Bempedoic acid (BA) and SB-204990 inhibit cyst growth of IMCD-derived *Pkd1*^*-/-*^ (ID1-3E5) kidney epithelial cells in 3D cultures. Cells were grown in Matrigel for a total of 7 days, supplemented with forskolin + IBMX after day 1, and then treated with either vehicle (DMSO), BA or SB-204990 for the last 3 days. **A**. Representative images of cystic structures under the different treatment conditions (scale bar = 100 µm). **B**. Summary data reveal that treatment with either BA or SB-204990 significantly reduced cyst area relative to CON (data from one experiment, n = 21-28, \*\*\*\**P* < 0.0001 for the indicated comparisons).

**Supplementary Figure 3.**
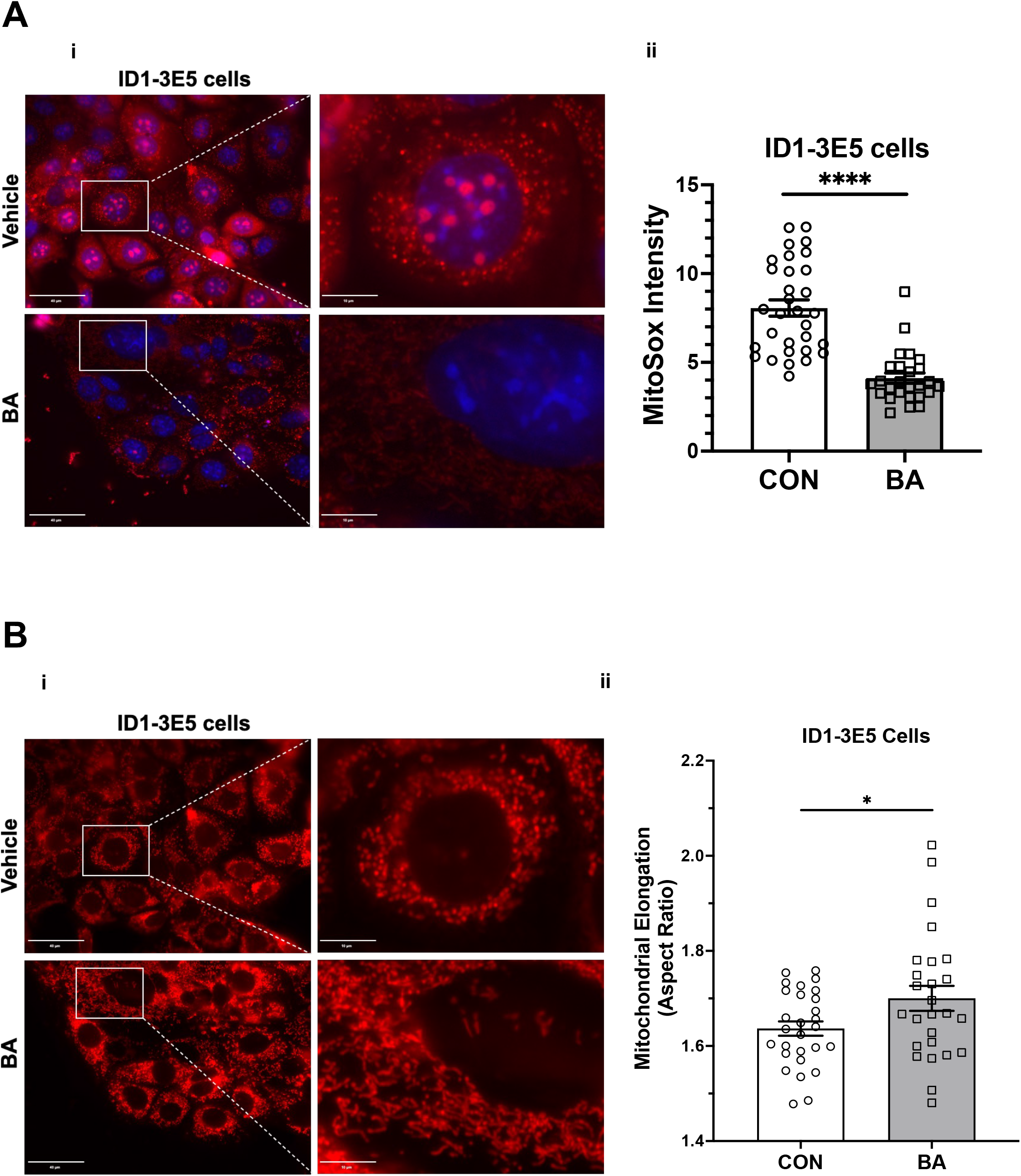
Bempedoic acid inhibits mitochondrial superoxide production and promotes mitochondrial elongation in IMCD-derived *Pkd1*^-/-^ kidney cells. **A**. To analyze the effect of BA on mitochondrial superoxide production, IMCD-derived *Pkd1*^*-/-*^ (ID1-3E5) cells were stained with MitoSOX™ Red mitochondrial superoxide indicator (*red*) and Hoechst 33342 nuclear stain (*blue*). **i**. Representative epifluorescence micrograph images are shown of ID1-3E5 cells in the absence (*top*) or presence (*bottom*) of BA treatment (100 µM) for 24 h. **ii**. Summary data reveal that BA treatment dramatically decreased mitochondrial superoxide production in ID1-3E5 cells (data from one experiment, n = 26-31 cells analyzed; \*\*\*\**P* < 0.0001). **B**. BA treatment significantly increased mitochondrial elongation of *Pkd1-*null cells. **i**. Representative images of MitoTracker Deep Red-stained ID1-3E5 cells in the absence (*top*) or presence (*bottom*) of BA treatment (100 µM) for 24 h. **ii**. Summary data reveal that BA treatment significantly increased mean cellular mitochondrial elongation in ID1-3E5 cells (mean cellular mitochondrial elongation values from one experiment with n = 26-28 cells analyzed, as described in *Materials and Methods*; \**P* < 0.05).

**Supplementary Figure 4.**
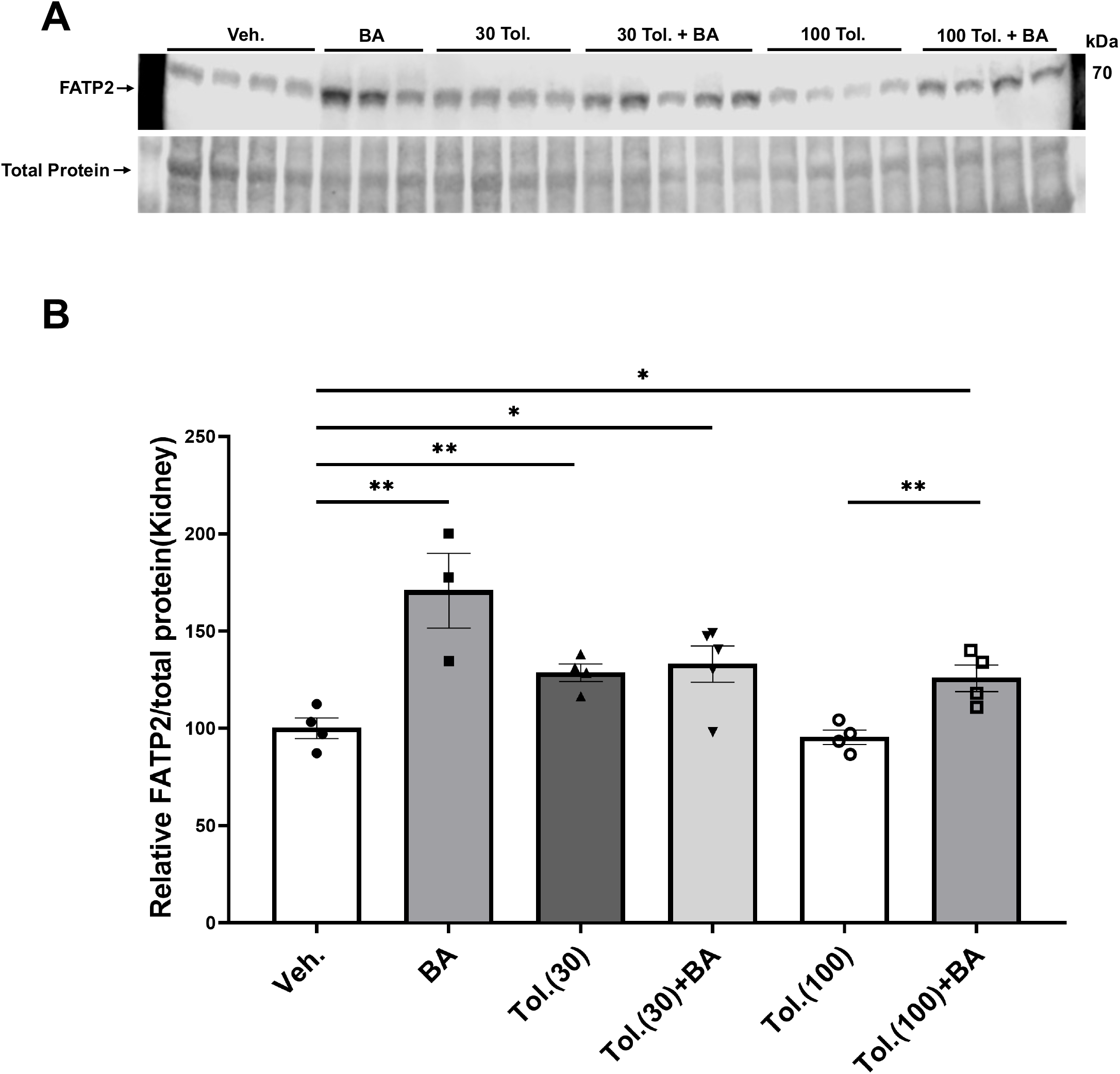
Effects of bempedoic acid and tolvaptan treatment in PKD mice on protein expression of the bempedoic acid-activating enzyme ACSVL1 (FATP2). **A**. *Upper*, immunoblot of kidney tissue homogenates from *Pkd1*^*fl/fl*^*;Pax8-rtTA;Tet-O-Cre* mice with doxycycline-induced *Pkd1* gene inactivation with or without concurrent treatment with BA (30 mg/kg/d) and/or tolvaptan (30 or 100 mg/kg/d) for FATP2a. *Lower*, total protein signal as loading control for the immunoblot. **B**. Densitometric quantifications of the FATP2a levels normalized to total protein indicate significant increases in BA- and tolvaptan-treated animals compared to vehicle control. (\**P* < 0.05, \*\**P* < 0.01 for the indicated comparisons).

